# Integrating lipid metabolism, pheromone production and perception by Fruitless and Hepatocyte nuclear factor 4

**DOI:** 10.1101/2023.02.23.529767

**Authors:** Jie Sun, Wen-Kan Liu, Calder Ellsworth, Qian Sun, Yu-Feng Pan, Yi-Chun Huang, Wu-Min Deng

## Abstract

Sexual attraction and perception, governed by separate genetic circuits in different organs, are crucial for mating and reproductive success, yet the mechanisms of how these two aspects are integrated remain unclear. In *Drosophila*, the male-specific isoform of Fruitless (Fru), Fru^M^, is known as a master neuro-regulator of innate courtship behavior to control perception of sex pheromones in sensory neurons. Here we show that the non-sex specific Fru isoform (Fru^COM^) is necessary for pheromone biosynthesis in hepatocyte-like oenocytes for sexual attraction. Loss of Fru^COM^ in oenocytes resulted in adults with reduced levels of the cuticular hydrocarbons (CHCs), including sex pheromones, and show altered sexual attraction and reduced cuticular hydrophobicity. We further identify *Hepatocyte nuclear factor 4* (*Hnf4*) as a key target of Fru^COM^ in directing fatty acid conversion to hydrocarbons in adult oenocytes. *fru*- and *Hnf4*-depletion disrupts lipid homeostasis, resulting in a novel sex-dimorphic CHC profile, which differs from *doublesex*- and *transformer*-dependent sexual dimorphism of the CHC profile. Thus, Fru couples pheromone perception and production in separate organs for precise coordination of chemosensory communication that ensures efficient mating behavior.

**Teaser:** Fruitless and lipid metabolism regulator HNF4 integrate pheromone biosynthesis and perception to ensure robust courtship behavior.

## Introduction

Chemical sensing, regarded from an evolutionary perspective as the oldest one, is common to all organisms, which are surrounded by a world full of odors emitted from conspecific or heterospecific individuals and the environment (*1*). Chemical communication is fundamental to social behaviors such as con-specific recognition, courtship, aggression, aggregation, and avoidance. Many animals, including insects, rely on chemical cues to locate and select appropriate mating partners for reproductive success. Insect chemical communication involves emission and perception of chemical cues (pheromones) and requires coordination of different organs that are involved in pheromone biosynthesis and sensing, respectively (*2*). *Drosophila*, with its long tradition in behavioral studies, is an ideal model to explore the evolution and diversity of pheromones associated with the diverse array of social behaviors, and the ethological context connecting chemical communication to behavioral and systemic processes. Although significant progress has been made in the *Drosophila* model during the past decades in unraveling the receptors and neural circuits for pheromone detection that give rise to innate behaviors, whether pheromone emission and perception can be regulated by the same upstream regulators remain unclear.

Insect cuticular hydrocarbons (CHCs), derived from long-chain fatty acids, are important for desiccation resistance (*3*–*5*), cold tolerance (*6*) and starvation resistance (*7*). Some CHCs function as sex pheromones for mate recognition (*8*–*12*). As in other insects, specialized hepatocyte-like oenocytes, located in the inner surface of the abdominal cuticle, are the primary site for biosynthesis of very long chain fatty acids (VLCFA) and VLCFA-derived hydrocarbons in *D. melanogaster* (*11*, *13*–*15*). Oenocytes are also the major site for ROS metabolism, proteasome-mediated protein catabolism, xenobiotic metabolism, ketogenesis, and peroxisomal beta-oxidation (*16*–*22*). Although the biosynthetic pathway for CHCs is active in both male and female oenocytes, the CHC profile shows apparent sexual dimorphism (*23*). In most populations of *D. melanogaster*, sexually dimorphic expression of enzymes such as *elongase F* (*eloF*) and *desaturase F* (*desatF*) leads to female-biased enrichment of several CHCs including 7,11-HD and 7,11-ND (*24*, *25*). Further proof that the CHC composition could be affected by sex-determination genes has been shown by ectopic expression of *transformer (tra*) in male oenocytes, which upregulates *eloF* and feminizes the CHC profile (*25*, *26*). However, the molecular mechanisms underlying the sexual dimorphism of CHC biosynthesis remain elusive (*27*).

Pheromone perception in *Drosophila* requires the neuronal function of *fruitless* (*fru*), a master regulator of the sexually dimorphic neural circuits that underlie sexual-dimorphic patterns of courtship (*28*–*31*), aggression (*32*–*34*), sleep (*35*) and other behaviors. The *fru* gene spans over 150 kb of the genome, and harbors P1–P4 promoters (*36*, *37*). The distal P1 promoter is dedicated to the expression of male-specific Fru proteins (Fru^M^), whereas P2–P4 promoters are involved in the production of non-sex-specific Fru proteins (Fru^COM^) (*37*–*39*) (Fig. 4A). Compared with the well-characterized involvement of Fru^M^ in neuronal behavior, the precise function of Fru^COM^ in development and behavior is less clear (*40*, *41*).

In this study, we report that Fru^COM^ is required for CHC biosynthesis in adult oenocytes of both sexes for chemical communication and desiccation resistance. Silencing of *fru* in oenocytes resulted in desiccation-sensitive adults with accompanying social behavioral changes. We further identify evolutionarily conserved Hepatocyte nuclear factor 4 (HNF4) as a key target of Fru^COM^ to inhibit steatosis and to direct fatty acid conversion to hydrocarbons in oenocytes. Thus, *fru* is involved in both pheromone biosynthesis and perception by distinct splicing isoforms in separate organs, bringing sexual attraction and perception under the control of a single gene.

## Results

### *fru* is required in both the nervous system and oenocytes for innate courtship behavior

To understand the function of Fru^COM^ isoforms, we induced whole-body clonal mutation of *fru* using a recently developed <u>m</u>osaic <u>a</u>nalysis by <u>g</u>uide (g) RNA-induced crossing-over (MAGIC) technique, which generates mosaic animals based on DNA double-strand breaks produced by CRISPR/Cas9 (*42*). The double gRNA design allowed us to eliminate all Fru isoforms in mutant clones randomly induced by a heat-shock inducible Cas9 (Fig. 4A). Interestingly, male flies bearing *fru* MAGIC clones, which were induced during early 3^rd^ larval instar, exhibited male-male courtship and male chaining behavior (Fig. 1A-D, and movie S1). This behavior is similar to Fru^M^ mutant flies reported previously (*43*, *44*), and the male-male courtship phenotype was reproduced when we used a pan-neuron *elav*-*Gal4* to knockdown *fru* expression (*elav*>*fru^IR^*) with a double-stranded (ds) RNA targeting exon C3, which is present in all Fru isoforms (Fig. 1A-D and 4A, movie S2).

**Fig. 1.**
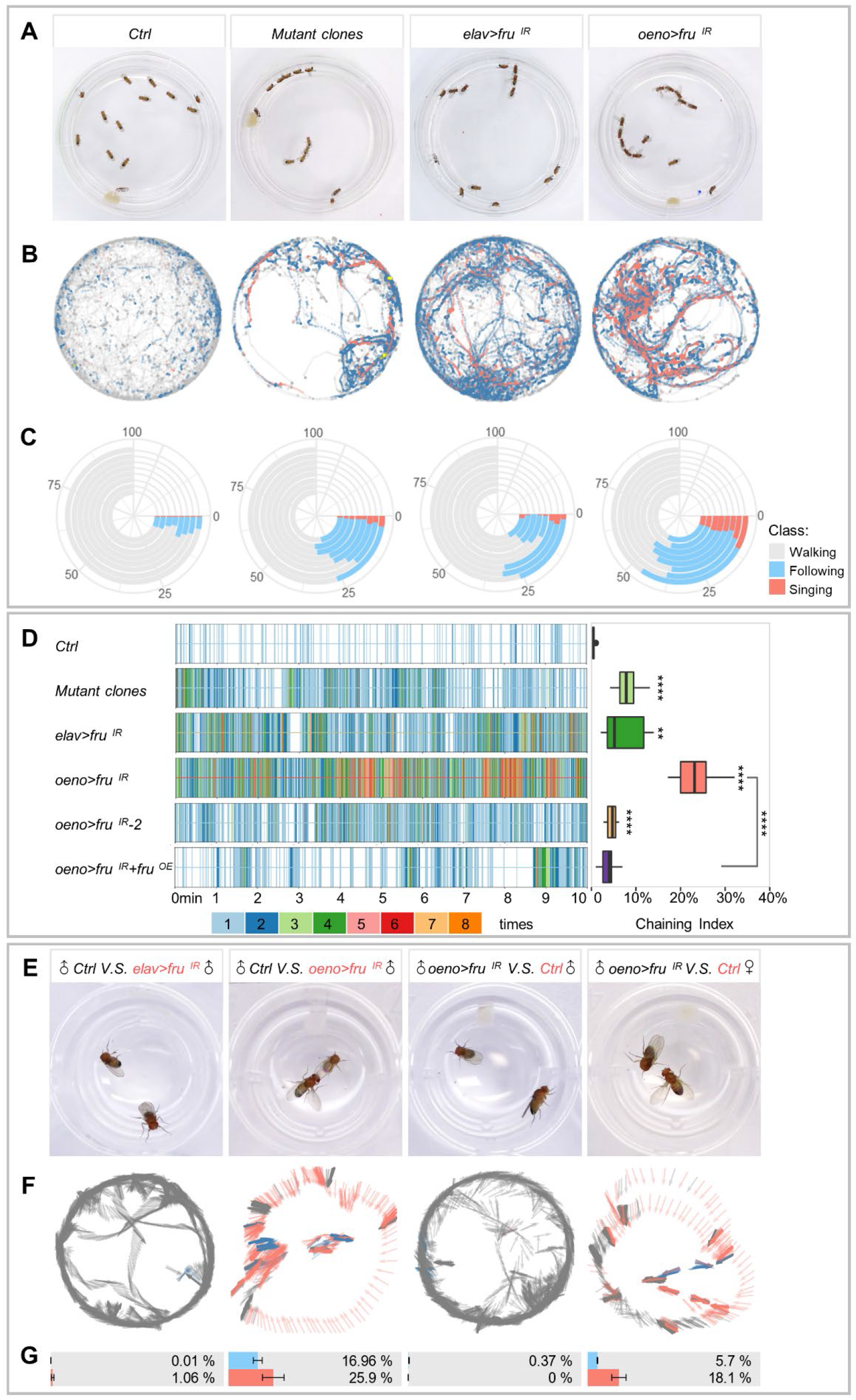
Loss of *fru* function in the nervous system or oenocytes alters male courtship behavior. (A) Representative movie screenshots of fly groups with indicated genotypes. 13 male adults of the same genotype were collected and placed in one chamber. (B) Event maps generated from representative movies of indicated genotypes (1.5-min movie per map). (C) Polarized bar plots illustrating the ratio of each detected behavior. Each bar plot was generated from 8 independent biological replicates (10-min movie per replicate). (D) Rug plots showing the chaining events detected from 10min movies. X axis refers to the time of movie. Each vertical line with different color stands for the number of chaining events detected in a single frame. The corresponding color to the number of events is illustrated in the legend. The box plots of chaining index are quantitative statistics of 8 independent biological replicates. (E) Representative movie screenshots of pair-housed flies with indicated genotypes (black: active party; red: passive beheaded party; *control*: *oeno*-*Gal4*/+) The trajectory and behavioral classification of the active party are shown in F. The arrow indicates the head orientation of the active party flies. The color of the arrow indicates different behavior status (grey: resting/walking; blue: following; salmon: singing) (1.5-min movie per map). Bar plots with error bar are quantitative statistics of 10 min movies of 8 independent biological replicates in G. Data are represented as mean ± SEM. P values are calculated using one-way ANOVA (D) and two-tailed unpaired t-test (G) followed by Holm-Sidak multiple comparisons. n.s., not significant, *p<0.05, **p<0.01, ***p<0.001, ****p<0.0001.

To better assay the behavior phenotype of different *fru* loss-of-function (LOF) flies, we developed a <u>m</u>achine-learning-based <u>a</u>utomatic <u>f</u>ly-behavioral <u>d</u>etection and <u>a</u>nnotation (MAFDA) system to track flies and identify multiple classes of behavior such as chasing, singing, and copulating (fig. S1A-E). The MAFDA platform facilitates visualization of complex behavioral phenotypes under a temporal and spatial overlay, and quantitative comparison of behavioral indexes among different genetic backgrounds. Using this platform, we found that the male-male following, singing and chaining behaviors exhibited in flies with MAGIC-induced *fru* mutant clones were comparable to or more pronounced than neuronal *fru* knockdown (Fig. 1D, and fig. S2A). The severe behavior phenotypes displayed by MAGIC *fru* mosaic flies were thus probably not due only to abnormalities in the nervous system, but also in the production of pheromones.

Oenocytes, which perform certain lipid processing functions, are the principal site for the synthesis of VLCFA-derived CHCs, including sex pheromones, that are associated with a variety of social behaviors in *Drosophila* (*11*, *13*, *14*). Using an oenocyte-specific Gal4 (*oeno-Gal4*) (*45*) to knockdown *fru* (*oeno*>*fru^IR^*) we found that male adult flies displayed male-male courtship and chaining behaviors (Fig. 1A-D, and movie S3). MAFDA analysis revealed that the male-male courtship behaviors exhibited by *oeno*>*fru^IR^* flies were highly pronounced (45% following, 25% chaining and 8% singing; Fig. 1D, and fig. S2A), and could be alleviated by Fru^COM^ overexpression in oenocytes (Fig. 1D, and fig. S2A, C), indicating that the defect was indeed caused by *fru* silencing. To avoid possible off-target effects of the dsRNA, additional independent *fru* RNAi lines were used and showed similar male-male courtship phenotypes (Fig. 1D, and fig. S2B). As revealed by MAFDA analysis, the degree of male-male courtship behavior by distinct RNAi lines was consistent with their respective knockdown efficiency (Fig. 1D and 4C, and fig. S2A). In addition, we combined gene-switch (GS) with *oeno*-*Gal4* to knockdown *fru* after pupal eclosion to avoid a possible role of *fru* in oenocyte development from its larval progenitors. Indeed, the behavioral phenotypes were similar between *fru*-depleted flies with or without gene-switch in oenocytes (fig. S2D). These results suggest that *fru* is required in oenocytes to maintain male courtship rituals.

To verify that the male-male courtship behavior in *oeno*>*fru^IR^* flies was caused by changes in sexual attraction rather than pheromone perception, we performed the single-pair mating experiment by placing a male fly with a headless target in the same chamber. The wild-type male was attracted to the headless male with *fru*-depletion in oenocytes (*oeno*>*fru^IR^*), but not the headless male lacking Fru in pan-neurons (*elav*>*fru^IR^* (Fig. 1E-G). By contrast, the male fly with oenocyte-specific *fru-* knockdown (*oeno*>*fru^IR^*) selected the decapitated wild-type female to initiate courtship but averted the wild-type male (Fig. 1E-G). These results indicate that while males lacking Fru in oenocytes maintain the ability in sexual selection, they are attractive to other males. Further, to determine whether females with oenocyte-depletion of Fru (*oeno*>*fru^IR^*) also show altered sexual attraction, we performed the two-choice courtship assay by putting a wild-type male and two headless female targets (*oeno*>*fru^IR^* and control *oeno*-*Gal4*/+) in the same chamber. MAFDA analysis revealed that the courtship index for *oeno*>*fru^IR^* females was ~60% of that of the control group (fig. S2E), suggesting that females with oenocyte *fru*-knockdown show reduced sexual attractiveness to male flies. Taken together, these results demonstrate that a deficiency of *fru* in oenocytes modifies sexual attraction in both sexes.

### Flies with *fru*-depletion in oenocytes exhibit aberrant social behavior

In *Drosophila*, innate social behaviors driven by pheromones include aggregation, male courtship, and female post-mating behaviors (*46*). To test whether *fru* knockdown affects social aggregation, we measured the distance between individual flies and their nearest neighbors: their ‘social space’ (*47*), within a social group. *oeno*>*fru^IR^* flies, on average, showed reduced social space and had more nearby neighbors when compared with the control (Fig. 2A), suggesting that *fru* expression in oenocytes is necessary for flies to maintain their social distance. Additionally, *oeno*>*fru^IR^* male flies showed reduced foraging behavior in a fixed time window (10 min) (Fig. 2B). MAFDA analysis revealed that the foraging index of the *oeno*>*fru^IR^* males was ~60% of the control group (Fig. 2B’’). Measurement of solid food intake in adult via a dye tracer further revealed that flies with *fru*-depletion ingested less food than wild-type flies over the same time period (1hr) (Fig. 2B’). To rule out that the observed social behavior changes are caused by impaired motor ability, we further examined their locomotion activity during the daytime. Compared with the control group, *oeno*>*fru^IR^* flies were more active, showing a 10% reduction in resting, a 35% increase in walking, and a 55% increase in leaping by MAFDA analysis (Fig. 2C), suggesting that the observed social behavior changes are not caused by impaired motor ability. Taken together, these data provide evidence that *fru* is required in oenocytes for normal social behavior.

**Fig. 2.**
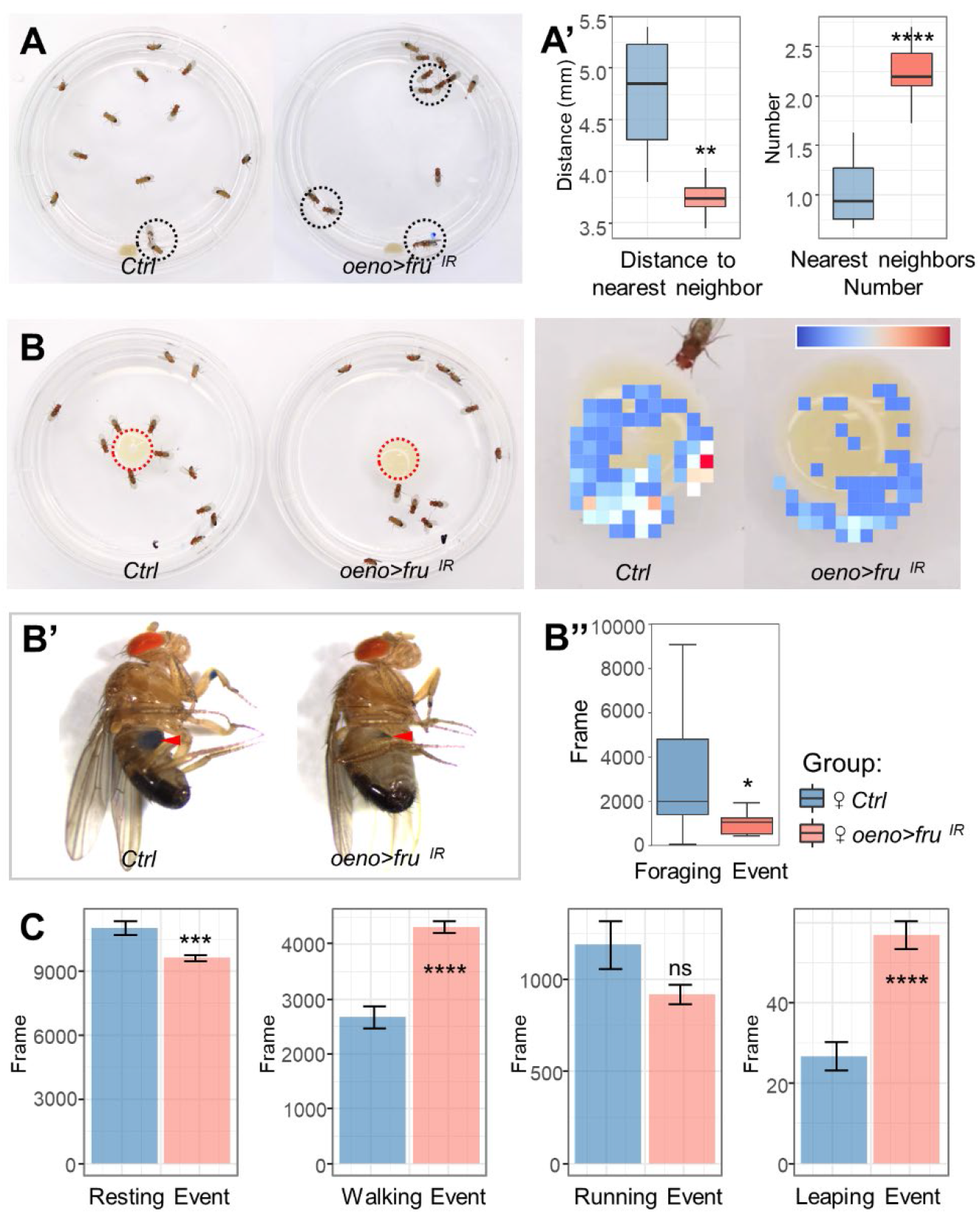
Social behavior changes in flies with *fru*-depletion in oenocytes. (A) Social space behavior. Left: representative images of an open field assay. Black circles indicate flies staying close together. (A’) Quantification of the distance of each fly to its nearest neighbor and the number of surrounding neighbors of each fly for both control (*oeno*-*Gal4*/+) and *oeno*>*fru^IR^* males (n = 8 groups of 13 flies for each genotype) (10-min movie per replicate). (B) Control (*oeno*-*Gal4*/+) and *oeno*>*fru^IR^* males were analyzed for total number of social interactions in a ‘competition-for-food’ assay. The heatmap on the right shows the degree to which flies gather in the food area. (B’) Measurement of solid food intake in adult using a dye tracer. (B’’) Quantification of the total number of feeding times of the flies for control (*oeno*-*Gal4*/+) and *oeno*>*fru^IR^* males (n = 10 groups of 13 flies for each genotype) (10-min movie per replicate). (C) Locomotion activity of control (*oeno*-*Gal4*/+) and *oeno*>*fru^IR^* males was monitored. From left to right, bar plots show resting events, walking events, running events and jumping events of flies, respectively (n = 8 independent experiments with 13 flies per genotype). For all paradigms, data are represented as mean ± SEM. P values are calculated using one-way ANOVA followed by Holm-Sidak multiple comparisons. Asterisks illustrate statistically significant differences between conditions. n.s., not significant, *p<0.05, **p<0.01, ***p<0.001, ****p<0.0001.

### *fru* function is necessary for CHC production

In *Drosophila*, CHCs act primarily as pheromones and play fundamental roles in sexual attraction or repulsion (*11*, *48*). Using gas chromatography-mass spectrometry (GC-MS) analysis, we identified 46 distinct chromatogram peaks from the cuticle extracts of wild-type adults and determined the chemical identities of their corresponding hydrocarbons (Fig. 3A, and fig. S3A-B). Subsequent analysis of CHC profiles by GC-MS and comparison of the main cuticular extracts of *elav*>*fru^IR^* and *oeno*>*fru^IR^* males revealed that the levels of total hydrocarbons were reduced by 90% when *fru* was knocked down in oenocytes, while *fru-loss* in the nervous system showed a CHC profile largely resembled the wild-type control (Fig. 3B and fig. S4A). Comparison of the CHC content in *oeno*>*fru^IR^* male flies revealed that 7-tricosene (7-T), the principle non-volatile male pheromone (*11*), was reduced by 97% (Fig. 3B’). Because 7-T mediates repulsion to other males and prevent male-male interactions (*45*, *49*), the drastic reduction of 7-T explains why wild-type males show a strong preference of males with *fru*-knockdown in the oenocyte (Fig. 1A-D). In addition, *oeno*>*fru^IR^* males showed a 87% decrease of *n*-tricosane (*n*C23), another characteristic pheromone of sexually mature fly which negatively regulates courtship and mating (*50*), may also underly the changed behavior patterns (Fig. 3B’).

**Fig. 3.**
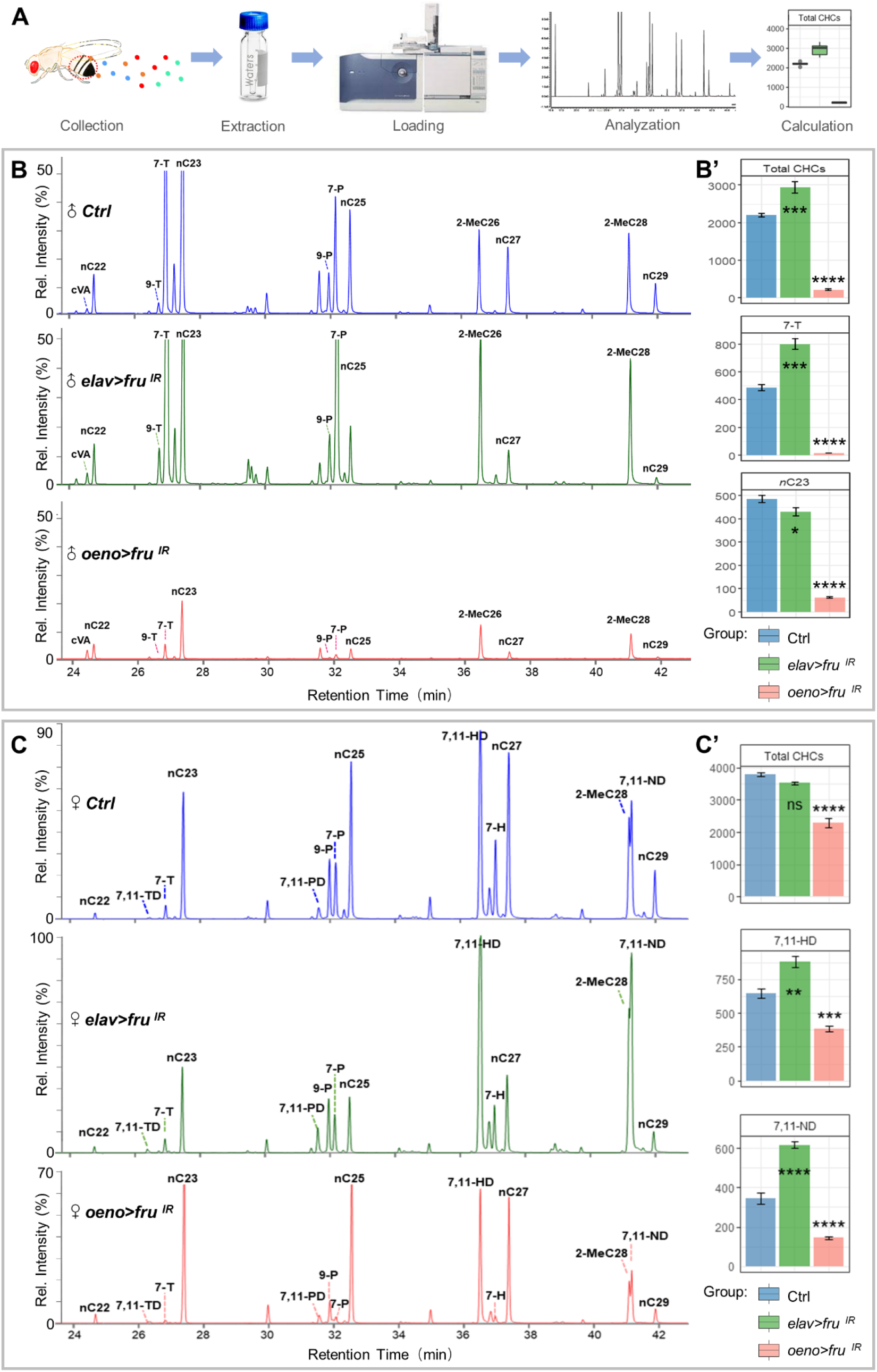
Fru is required in the oenocytes for the biosynthesis of cuticular hydrocarbons. (A) A schematic of cuticular hydrocarbon extraction and GC-MS analysis. (B) CHCs from adult male flies of each genotype were analyzed using GC-MS. Compared with the *control* and *elav*>*fru^IR^* males, *oeno*>*fru^IR^* males exhibit significantly lower levels of CHCs. (B’) The absolute contents of total hydrocarbons, male key pheromones (7-T and nC23) carried by a single male were calculated by the loading of internal standards. (C) CHCs from females of each genotype were analyzed using GC-MS. Compared to *controls* and *elav*>*fru^IR^* females, *oeno*>*fru^IR^* females exhibit lower levels of CHCs. (C’) The absolute contents of total hydrocarbons, female key pheromones (7,11-HD and 7,11-ND) carried by a single female were calculated by the loading of internal standards. Data are represented as mean ± SEM. P values are calculated using one-way ANOVA followed by Holm-Sidak multiple comparisons. Asterisks illustrate statistically significant differences between conditions. n.s., not significant, *p<0.05, **p<0.01, ***p<0.001, ****p<0.0001.

CHCs are highly sexually dimorphic in *D. melanogaster*, with many of the compounds present in one sex but absent in the other, while shared compounds often differ between sexes (*51*). To determine whether *fru* knockdown also alters the CHC profile in female flies, we performed the GS-MS analysis and found that the total CHCs were reduced by 40% in *oeno*>*fru^IR^* females when compared with the control. Different from the male flies, the changes in the CHC profile fall mainly into the CHCs with the chain length beyond C26 (Fig. 3C), which include the female pheromones 7,11-HD and 7,11-ND (*45*). These results explain why wild-type female are more sexually attractive than females with *fru-*knockdown in oenocytes (fig. S2E).

The hydrophobic properties of CHCs protect insects from transpirational water loss through insect epicuticle and play a major role in desiccation resistance (*14*, *52*). In *oeno*>*fru^IR^* flies, we found significant reduction of multiple long-chain *n*-alkanes, including *n*-heptacosane (*n*C27) and *n*-nonacosane (*n*C29) (Fig. 3B-C and fig. S4A). To test the hydrophobicity of these flies, both male and female adults were dehydrated and incubated in water to assess defects in hydrophobic coating of the cuticle. While control carcasses remained at the surface of the liquid following a 4-hr incubation, *oeno*>*fru^IR^* carcasses sank below the surface (fig. S4B-C). Together, these results suggest that *fru* is required in both male and female oenocytes for CHC and sex pheromone production.

### Fru^COM^, not Fru^M^, is expressed in the oenocytes

*fru* is a complex locus with sophisticated precise spatiotemporal control of transcription through four distinct promoters (*36*, *37*, *41*) (Fig. 4A). Translation of P1 transcripts in males produces the male-specific Fru^M^ proteins that have an amino-terminal extension of 101 amino acids preceding the BTB domain, whereas transcripts from the P2–P4 promoters encode a set of non-sex-specific Fru^COM^ proteins that have essential functions in the development of both sexes (*38*) (Fig. 4A). All Fru proteins are putative transcription factors containing a common BTB N-terminal domain and five alternatively spliced C-terminus varying in the number and sequence of Zn-finger DNA binding domains (A, B, C, D, or E) (*36*, *37*, *53*) (Fig. 4A).

**Fig. 4.**
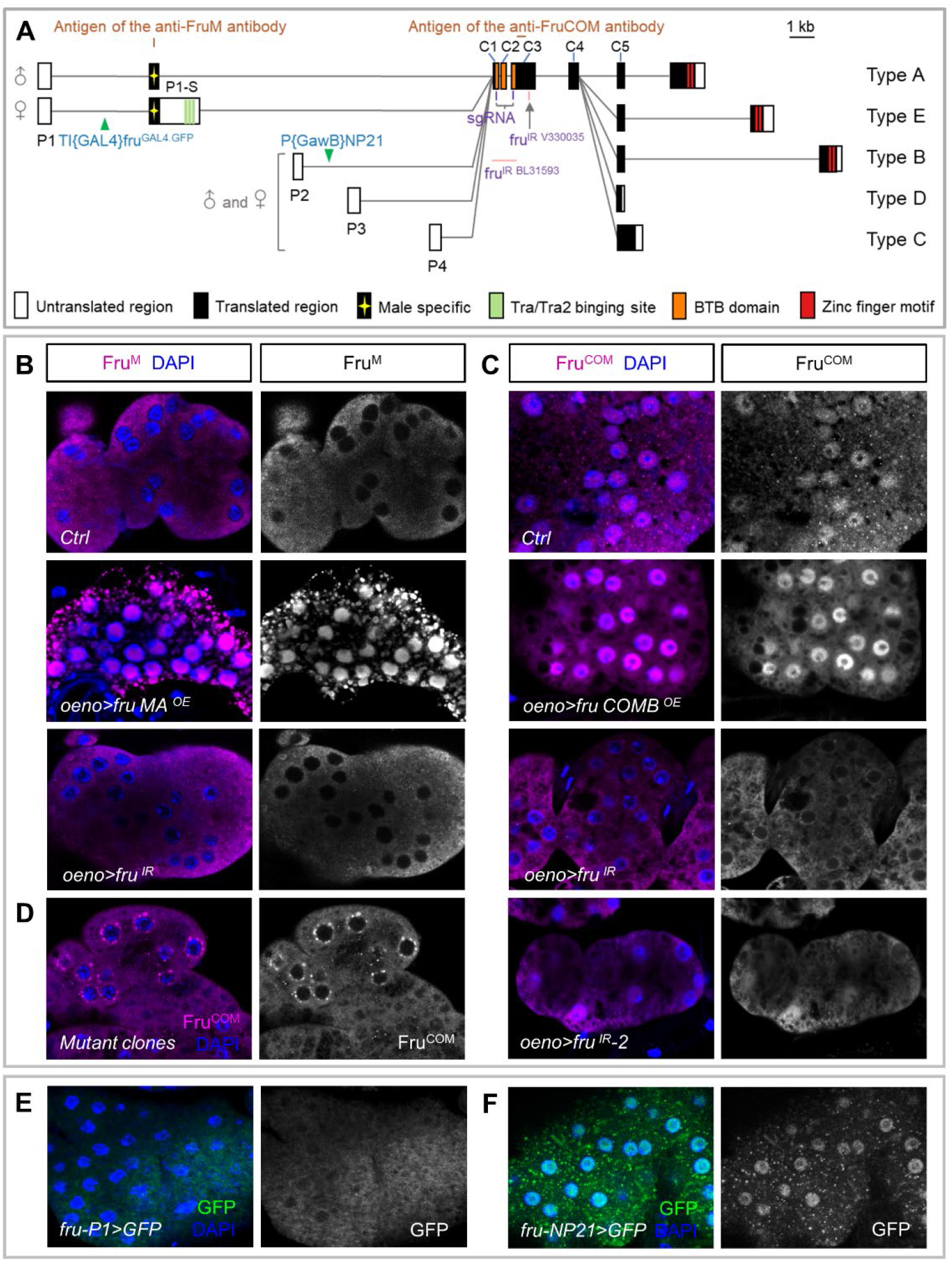
The *fru* gene locus and its expression in oenocytes. (A) Schematic representation of the *fru* gene locus. Locations of four promoters (P1–P4), the exonintron organization, and the P-element insertion sites of *fru^P1^* and *fru^NP21^* (green triangles) are shown. Filled and open boxes indicate coding and non-coding exons, respectively. A-E denote isoform-specific exons for types A-E. The start and termination codons are also shown. The regions containing epitopes for the anti-Fru antibodies are indicated. *fru* gRNA and *fru* dsRNA sites are displayed. (B and C) Anti-Fru antibodies were used to stain oenocytes, from wild-type flies and overexpression of Fru^M^/Fru^COM^ or knockdown of Fru (all isoforms) in oenocytes. (D) Validation of the effectiveness of *fru* mutant clones by anti-Fru^COM^ staining in oenocytes. (E and F) *fru*-*P1*-*Gal4* inserted into the first intron and *fru-NP21-Gal4* inserted into the second intron of *fru*, driving *UAS*-*GFP* as a reporter to mark the transcriptional level of *fru* in oenocytes.

To determine which Fru isoforms are expressed in oenocytes, we stained oenocytes with a Fru^M^-specific antibody (*54*) and found no signal (Fig. 4B). Using a newly generated polyclonal antibody recognizing an epitope in exon C3, which is present in both Fru^COM^ and Fru^M^ isoforms (Fig. 4A), we detected strong signal in oenocyte nuclei in both male and female adults (Fig. 4C). The Fru^COM^ signal was lost in oenocytes with *fru RNAi* knockdown or with MAGIC induced *fru* deletion (Fig. 4C-D). Furthermore, we found that *fru*-*P1*-*Gal4* (*55*), with the Gal4 inserted in the first intron to mimic the Fru^M^ pattern (Fig. 4A), showed no expression in oenocytes (Fig. 4E). In contrast, *fru-NP21-Gal4*, with its Gal4 located in the second intron (*56*), possibly under the control of both P1 and P2 promoters (Fig. 4A), drove UAS-GFP expression in oenocytes (Fig. 4F). Together, these results suggest that Fru^COM^, not Fru^M^, is expressed in oenocytes in both sexes to regulate CHC biosynthesis.

### Fru^COM^ controls *Hnf4* expression in oenocytes to maintain lipid homeostasis

In insects, long-chain fatty acid (LCFA) biosynthesis is predominantly derived from palmitate, which is synthesized by *Acetyl*-*CoA carboxylase* (*ACC*) and *Fatty acid synthase 2* (*FASN2*) (*57*). After that, they are elongated by the successive action of elongases, desaturases and decarbonylate enzymes, before being converted to hydrocarbons (*14*, *52*, *58*). To determine how Fru^COM^ affects hydrocarbon biosynthesis, we generated multiple RNA-Seq datasets from male oenocytes with *fru*-knockdown by three independent *fru*-RNAi lines and corresponding control samples, which were then subjected to Principal Component Analysis (PCA). On the PCA plot, the three independent *fru*-RNAi datasets clustered together, and separate from the control samples (fig. S5A-B). When compared with the wild-type control, we found that the transcripts of genes involved in various steps of VLCFA and hydrocarbon biosynthesis, including *ACC*, *Hepatocyte nuclear factor 4* (*Hnf4*), *FASN2*, *Desaturase1* (*desat1*), *Cytochrome P450 4g1* (*Cyp4g1*), and multiple *elongases* (*CG16904* and *CG30008*), were reduced in oenocytes with *fru*-knockdown (Fig.5A). The downregulation of these transcripts was validated by RT-qPCR of RNAs from male oenocytes (Fig. 5B).

**Fig. 5.**
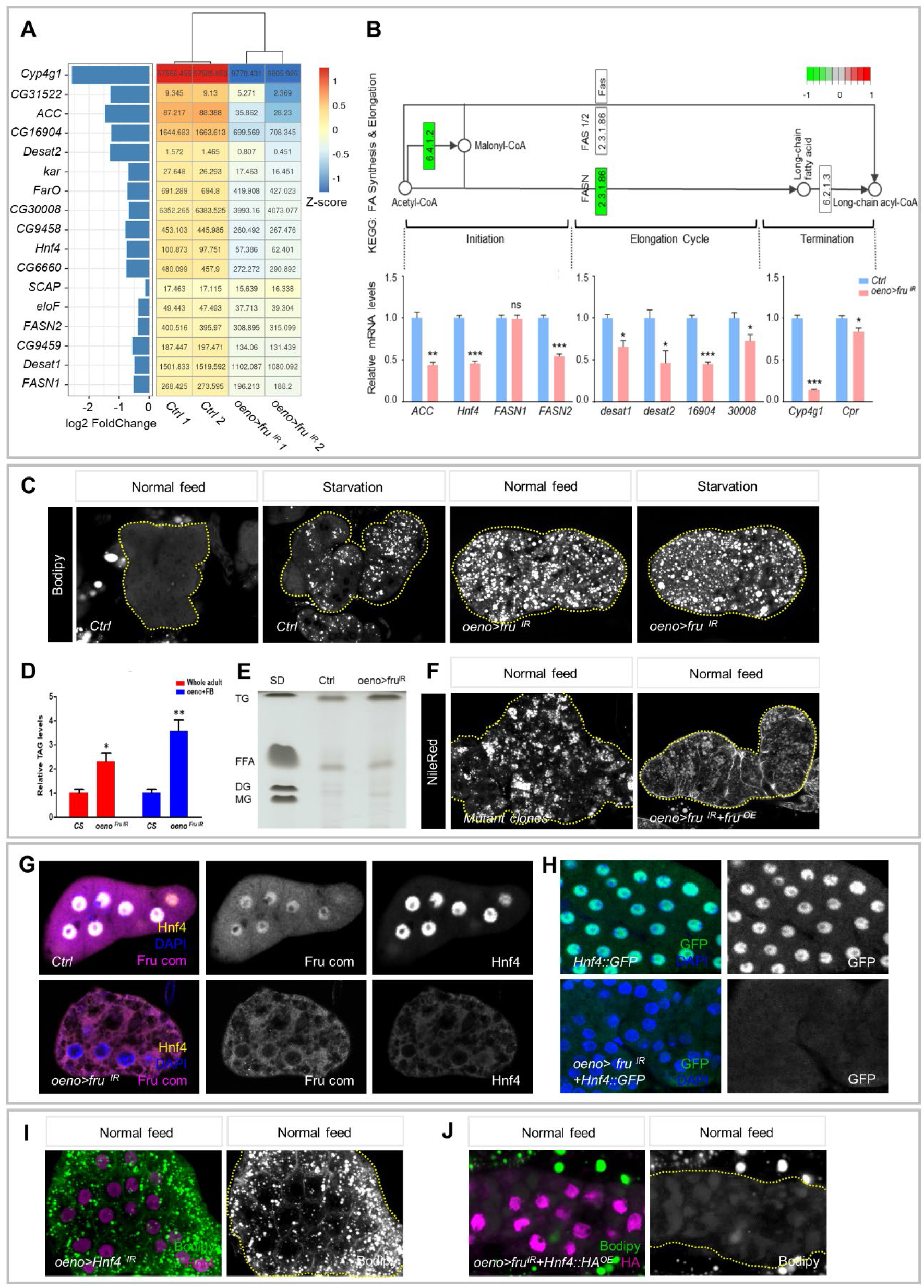
Fru controls HNF4 expression in the oenocytes to inhibit steatosis. (A) Downregulation of VLCFA/hydrocarbon biosynthesis genes in oenocytes with *fru*-knockdown from two independent replicates. Colors in the heatmap correspond to the scaled FPKM which is shown as a number in each cell. (B) The FA synthesis & elongation pathway from KEGG, with green boxes indicating genes down-regulated in this subset, and RT-qPCR analysis of genes in the VLCFA/hydrocarbon metabolic pathway (controls: blue bars; *oeno*>*fru^IR^*: salmon bars). Transcript levels are normalized to *Rp49* mRNA and presented relative to the level of controls. Asterisks illustrate statistically significant differences between conditions. n.s., not significant, *p<0.05, **p<0.01, ***p<0.001. (C) Bodipy stains are depicted for oenocytes dissected from control and *oeno*>*fru^IR^* males raised on standard diet and starved for 7-day after emergence. Oenocytes are outlined with a yellow dotted line. (D) Triglyceride levels were measured in 7-day old control and *oeno*>*fru^IR^* reared on a standard diet after emergence. Metabolite levels are normalized to total protein and presented relative to the amount in control animals. Red indicates that decapitated whole body was used as the sample material, and blue indicates that only adipose tissue (fat body and oenocytes) was used. (E) TLC analysis shows TAG and FFA levels in adipose tissue of control and *oeno*>*fru^IR^* males. (F) NileRed stains are depicted for oenocytes dissected from *fru mutant clones* and *oeno*>*fru^IR^*+*fru^OE^* males cultured on standard diet. (G) *fru*-knockdown in oenocytes resulted in decreased HNF4 protein levels. Upper panel, Fru (magenta) colocalized with HNF4 (yellow) in control oenocytes. Lower panel, the level of HNF4 was significantly reduced in oenocytes of *fru*-knockdown. (H) GFP-tagged HNF4 from the endogenous *Hnf4* promoter is highly expressed in oenocytes, but the GFP signal is lost in oenocytes with *fru*-knockdown. (I) Bodipy (green) and anti-HNF4 (magenta) stain are depicted for oenocytes dissected from *oeno*>*Hnf4^IR^* males raised on standard diet. (J) Bodipy stains are depicted for oenocytes of *oeno*>*fru*^*IR*+^*Hnf4^OE^* males cultured on standard diet.

The hepatocyte-like oenocytes accumulate lipid droplets (LDs) as a normal response to fasting and this readout of steatosis also indicates abnormal lipid metabolism in fed flies (*17*, *45*). We noticed LD accumulation in oenocytes in both starved and fed *oeno*>*fru* ^*IR*^ flies, suggesting that *fru* is required for preventing steatosis independent of nutritional status (Fig. 5C). To determine whether *fru-* knockdown interferes with systemic lipid homeostasis, we measured the TAG levels in *oeno*>*fru* ^*IR*^ male adults and found that the whole-body TAG level of these flies was 2.4-fold of the control (Fig. 5D), a result that explains why *fru*-knockdown males have less food requirements than control males (Fig. 2B). In addition, thin layer chromatography (TLC) of adipose tissue revealed that the TAG content, but not free fatty acids (FFA), was significantly increased when *fru* was depleted in oenocytes (Fig. 5E). Consistently, flies bearing *fru* MAGIC clones exhibited similar LDs accumulation, and *fru*-depletion induced steatosis could be alleviated by Fru^COM^ overexpression in oenocytes (Fig. 5F). These results indicate that Fru functions in oenocytes to prevent steatosis and to maintain systemic lipid homeostasis.

Among the downregulated genes induced by *fru*-depletion, we focused on *Hnf4*, which has been reported to be necessary in maintaining lipid homeostasis and promoting fatty acid conversion to VLCFA and hydrophobic hydrocarbons in oenocytes (*59*). To determine whether HNF4 acts downstream of Fru, we first used an anti-HNF4 antibody (*60*) and found that the HNF4 protein level was significantly downregulated in *fru*-silenced oenocytes (Fig. 5G, and fig. S6A). Consistently, a HNF4 protein trap line with GFP-tagged endogenous HNF4 (*61*), showed loss of the GFP signal in oenocytes with *fru*-depletion (Fig. 5H), further suggesting that HNF4 expression is regulated by *fru* in oenocytes. To determine whether *Hnf4* is functionally downstream of *fru* in oenocytes, the dsRNAs of *Hnf4 (Hnf4^IR^*) were targeted to oenocytes by *oeno-Gal4*. As expected, knockdown of *Hnf4* in oenocytes resulted in similar steatosis, as reported previously (*59*), whereas HNF4 overexpression alleviated lipid accumulation induced by *fru*-depletion (Fig. 5I-J). Taken together, these results suggest that Fru^COM^ regulates HNF4 expression in oenocytes to inhibit steatosis and maintain systemic lipid homeostasis.

### *Hnf4* is essential in mediating *fru*-regulated CHC biosynthesis and courtship behavior

To determine whether *fru*-regulated CHC production is mediated by HNF4, we used GC-MS to measure hydrocarbon levels in flies with HNF4 depletion in oenocytes (*oeno*>*Hnf4^IR^*). Unsurprisingly, these flies also show significant decreases in total CHCs in both sexes (94% decrease in males and a 62% decrease in females), similar to flies with *fru*-knockdown in oenocytes (Fig. 6A-B). Notably, knockdown of *Hnf4* also resulted in defective synthesis of sex pheromones in both sexes, including male pheromone 7-T (99% decrease), *n*C23 (89% decrease) and female pheromone 7,11-HD (66% decrease), 7,11-ND (80% decrease) (Fig. 6A-B). In addition, *oeno*>*Hnf4^IR^* flies also showed a significant reduction of multiple long-chain *n*-alkanes (Fig. 6A-B), resulting in defects in the hydrophobic coating of cuticles in both male and female adults (fig. S6B). Next, we asked whether re-introducing HNF4 in oenocytes with *fru*-knockdown would restore the CHC profile, and found that the total CHC levels were restored to 71% and 79% of the control males and females, respectively. It is noteworthy that the major sex pheromones in both male and female flies, especially 7-T and 7,11-HD, were well recovered and comparable to the levels of the control flies (Fig. 6A-B).

**Fig. 6.**
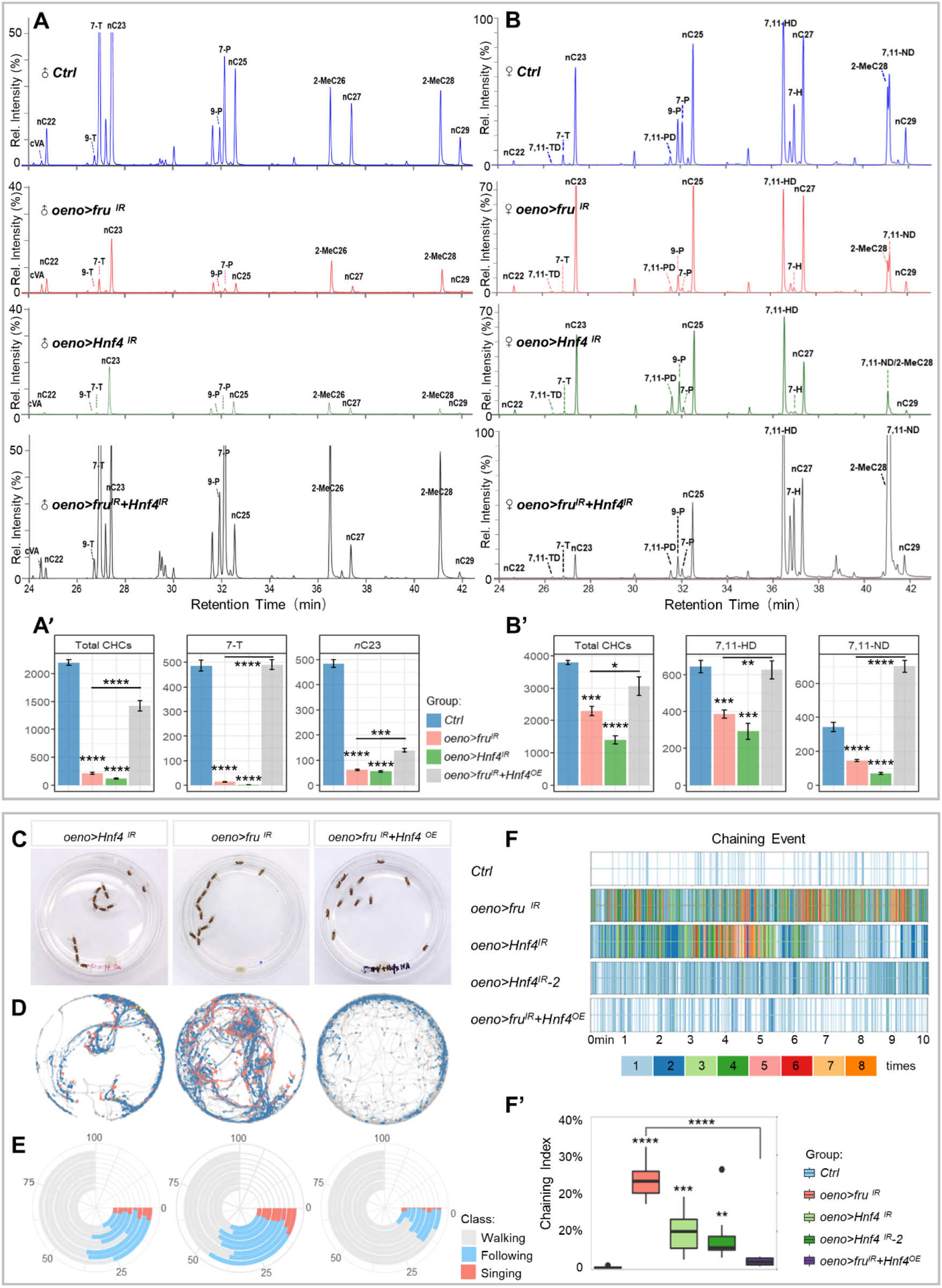
Fru controls HNF4 expression in the oenocytes to maintain CHC biosynthesis and innate courtship behavior. (A) GC-MS analysis of CHCs from male adults in *oeno*>*Hnf4^IR^* and *oeno*>*fru^IR^*+*Hnf4^OE^*. (A’) The absolute contents of total hydrocarbons, male key pheromones (7-T and nC23) carried by a single male were calculated by the loading of internal standards. (B) GC-MS analysis of CHCs from female adults of *controls* (*oeno*-*Gal4*/+), *oeno*>*fru^IR^*, *oeno*>*Hnf4^IR^* and *oeno*>*fru^IR^*+*Hnf4^OE^* females. (B’) The absolute contents of total hydrocarbons, female key pheromones (7,11-HD and 7,11-ND) carried by a single female were calculated by the loading of internal standards. Data are represented as mean ± SEM. P values are calculated using one-way ANOVA followed by Holm-Sidak multiple comparisons. Asterisks illustrate statistically significant differences between conditions. n.s., not significant, *p<0.05, **p<0.01, ***p<0.001, ****p<0.0001. (C) Representative movie screenshots of fly groups with indicated genotypes. 13 male adults of the same genotype were collected and placed in one chamber. (D) Event maps generated from representative movies of indicated genotypes (1.5-min movie per map). (E) Polarized bar plots illustrating the ratio of each detected behavior. Each bar plot was produced from 8 independent biological replicates (10-min movie per replicate). (F) Rug plots showing the chaining events detected from 10-min movies. X axis refers to the time of movie. Each vertical line with different color indicates the number of chaining events detected in a single frame. The corresponding color to the number of events is illustrated in the legend. (F’) The box plots of chaining index are quantitative statistics of 8 independent biological replicates.

The impairment of CHC biosynthesis suggests that flies with *Hnf4*-knockdown in oenocytes may also have changed courtship-behavior patterns. To test this, we applied the MAFDA platform to analyze the courtship behavior of male flies with oenocyte *Hnf4*-knockdown using two independent *Hnf4*-*RNAi* lines. As expected, these flies exhibited pervasive male-male courtship and chaining behaviors, similar to *fru*-depleted males (Fig. 6C-F, and fig. S6C-D and movie S4). Importantly, misexpression of HNF4 in oenocytes with *fru*-knockdown alleviated the male-male courtship and male-chaining phenotypes when compared with *fru*-knockdown alone (Fig. 6C-F, and fig. S6D and ovie S5), highly consistent with their respective CHC profiles (Fig. 6A). Together, these results suggest HNF4 is a key factor downstream of Fru in oenocytes to control pheromone and CHC production and its downregulation is accountable for the behavior phenotypes displayed by *oeno*>*fru^IR^* flies.

### *dsx*- and *fru*-depletion induce two distinct sex-dimorphic CHC profiles

Although *fru*- or *Hnf4*-knockdowns in oenocytes induced significant decreases of total CHCs in both male and female flies, the changes appeared sexually dimorphic. While males showed almost complete loss of CHCs, the reduction of CHCs in females was mostly on those with a chain-length beyond C26 (Fig. 3B-C). In insects, the sexual dimorphism of CHCs could be regulated by sex determination genes, including *tra* and *dsx* (*24*, *62*–*64*). Consistently, knockdown of *dsx* in oenocytes caused male-chaining behavior (Fig. 7A). To test whether *fru* acts downstream of *dsx* in oenocytes, we first performed immunohistochemistry analyses using anti-HNF4 and anti-Fru^COM^ antibodies and found that the protein levels of HNF4 and Fru^COM^ were not changed in oenocytes with *dsx*-knockdown (Fig. 7B). Further CHC profiling revealed that oenocyte-specific depletion of *dsx* led to a 26% increase in total CHCs in males but a 19% decrease in females (Fig. 7C-D). Consistent with these findings, *oeno*>*dsx^IR^* male flies showed better water repellency than controls, whereas females had impaired hydrophobic coating of the cuticle (Fig. 7A-B). And at the pheromone level, *dsx*-loss in oenocytes caused feminization of the pheromone profile in male flies and masculinization of the pheromone profile in females (Fig. 7C-D). These findings reveal that despite the phenotypic similarity in courtship behavior between *dsx*- and *fru*-deleted flies, the underlying mechanism of these two genes in regulating CHC and pheromone profiles is different (Fig. 7C-D). The mechanism underlying the sex differences of CHC profile in *fru* and *Hnf4* depleted flies is yet to be determined. Nonetheless, our results suggest that *fru* is not a downstream target of *dsx* in regulating CHC biosynthesis in oenocytes.

**Fig. 7.**
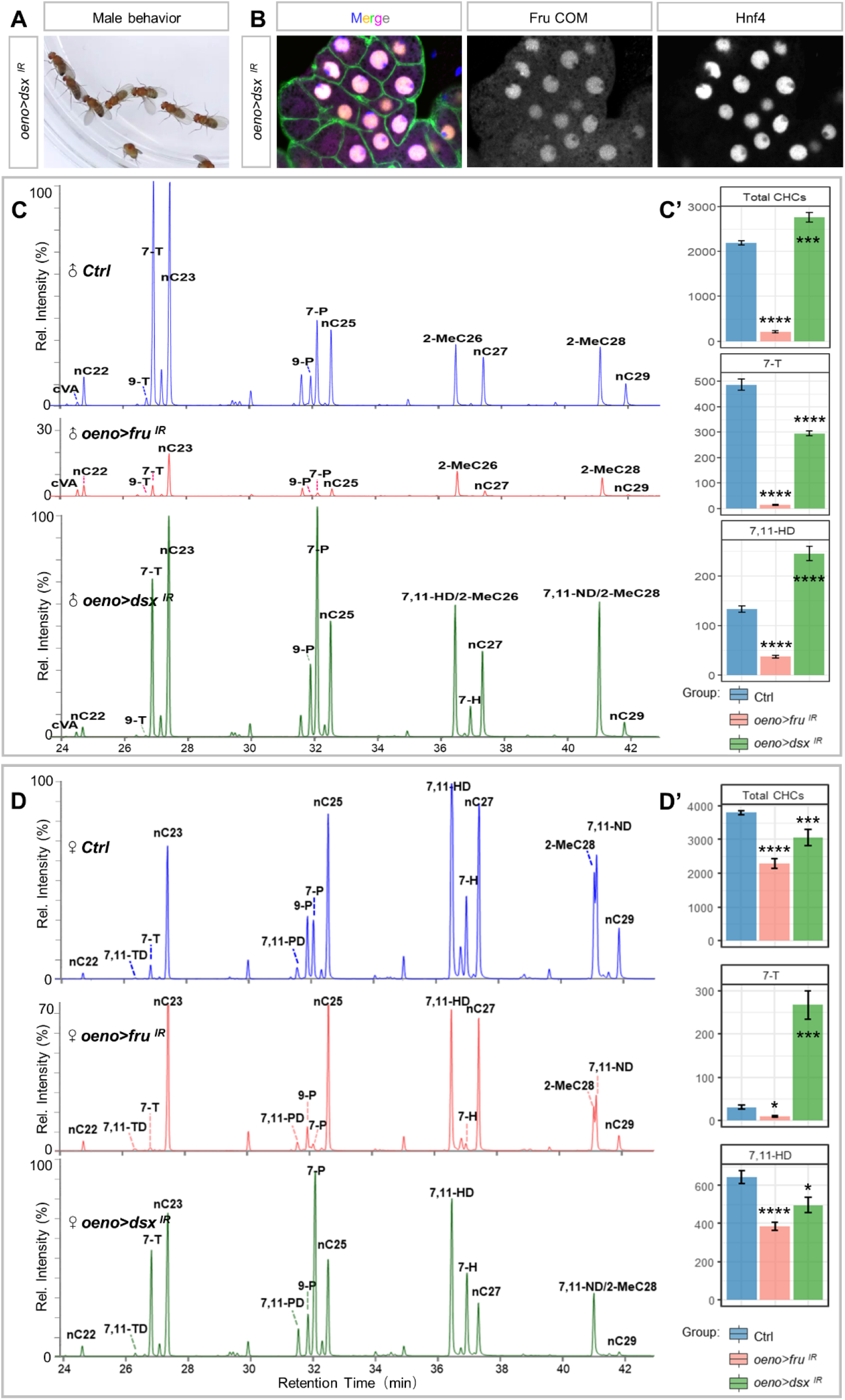
Fru does not act downstream of Dsx in regulating CHC biosynthesis. (A) Silencing of of *dsx* in oenocytes causes male chaining behavior. (B) Immunostaining shows that Fru^COM^ and HNF4 protein levels remain high in oenocytes with *dsx-*-knockdown. (C) CHCs from males of each genotype were analyzed using GC-MS. *oeno*>*fru^IR^* males exhibit significantly lower levels of CHCs in the full spectrum than the controls. The CHCs of *oeno*>*dsx^IR^* males, in contrast, exhibit a mixture of low-levels of characteristic male hydrocarbons (7-T) and high-levels of diene hydrocarbons (7,11-HD and 7,11-ND, characteristic of females). (D) GC-MS analysis of CHCs in female adults shows that oenocyte-specific knockdown (*oeno*>*fru^IR^*) resulted in lower levels of CHCs in the full spectrum than control females. However, knockdown of dsx in female oenocytes (*oeno*>*dsx^IR^*) resulted in lower female pheromones (diene hydrocarbons 7,11-HD and 7,11-ND) but high levels of male pheromone (7-T). Data are represented as mean ± SEM. P values are calculated using one-way ANOVA followed by Holm-Sidak multiple comparisons. Asterisks illustrate statistically significant differences between conditions. n.s., not significant, *p<0.05, **p<0.01, ***p<0.001, ****p<0.0001.

## Discussion

This study reveals a combined control of pheromone production and perception by a zinc-finger transcription factor gene, *fru*, which regulates lipid homeostasis through an evolutionarily conserved gene *Hnf4*, thereby promoting the robustness and efficiency of courtship behavior in *Drosophila*. Such an integrated regulation of sexual attractiveness and sexual perception/execution by a single gene in distinct cells may have reproductive advantage, as the recruitment of *fru* in certain insect species (*65*) could enhance both the emission and perception of sex-related cues simultaneously, which has stronger selective advantage than separate evolvement of each process (Fig.8).

**Fig. 8.**
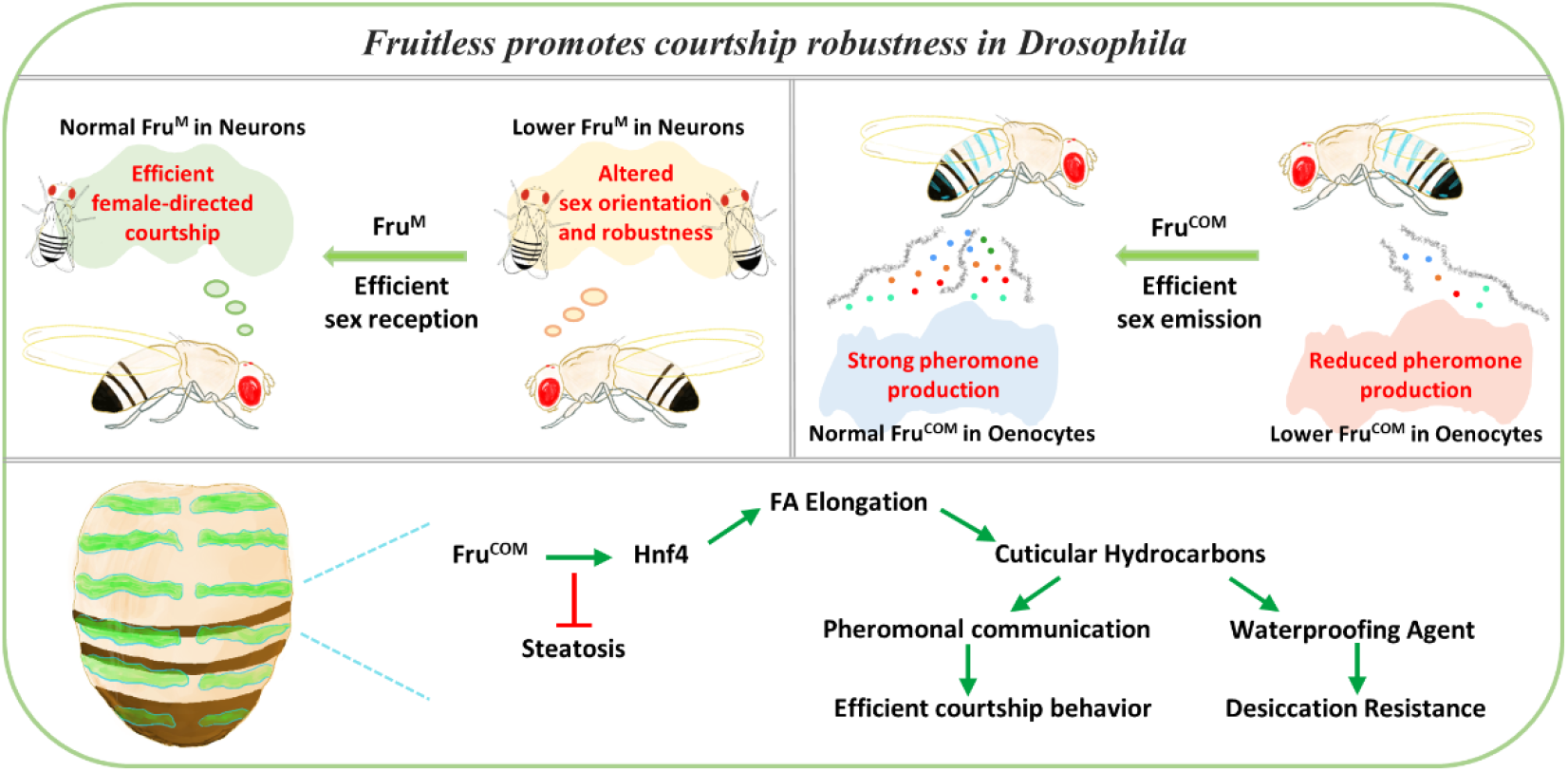
Model of *fruitless* regulates pheromone biosynthesis and perception. A schematic drawing to show that Fru regulates both pheromone production and perception through different isoforms (the male-specific Fru^M^ and the non-sex-specific Fru^COM^) expressed in different organs, thereby promoting the robustness and efficiency of courtship behavior. Fru^COM^ expression in oenocytes regulates HNF4 protein levels for the biosynthesis of sex pheromones, along with other cuticle hydrocarbons required for desiccation resistance.

The *fru* gene locus contains a complex transcription unit with multiple promoters and alternative splicing isoforms (Fig. 4A). *fru* P1 transcripts have only been detected in the nervous system, their sex-specific protein product Fru^M^ is expressed in ~2000 neurons to masculinize their structure and function (*66*). In contrast, Fru^COM^ is expressed in both neural and non-neural tissues. Its expression has been detected in neuroblasts in both male and female larval CNS (*41*). Fru^COM^ appears to be also required in adult female brain to regulate female rejection behavior (*67*). Outside the nervous system, Fru^COM^ is detected in specific cell types in the reproductive system (*66*, *68*), and is necessary for the maintenance of germline stem cells and cyst stem cells in male gonads (*40*). Our study shows that in pheromone-producing oenocytes, Fru^COM^, rather than Fru^M^, is expressed in both sexes and plays a key role in CHC biosynthesis (Fig. 4C). Although how the sex- and tissue-dynamic expression of Fru^M^ and Fru^COM^ is orchestrated remains to be resolved, the utilization of different transcriptional/splicing products of the same gene in pheromone production and perception may provide a potential advantage in coordinating these two related biological processes during development.

In insect, pheromone synthesis has coevolved with chemosensory perception. Numerous examples of correlated evolutionary changes between pheromone production and perception have been reported, with evolutionary pros and cons of sex-specific pheromones mirroring their sensory responses (*69*, *70*). The genetic mechanisms that control pheromone perception is best understood in *Drosophila* with >150 chemoreceptor proteins identified (*11*). The odorant receptor neurons (ORNs) that express ORs are specialized to detect most volatile chemicals, including low volatility pheromones (*71*). In Or47b ORNs, *fru* has been shown to regulate sensory plasticity and act as a downstream genomic coincidence detector (*72*–*75*). Although it is unclear whether *Hnf4* is regulated by *fru* in the nervous system, neuronal function of *Hnf4* has been reported during neural stem cell differentiation and in the aging brain for β-oxidation of fatty acids (*76*, *77*). Impaired lipid homeostasis has been shown to be associated with disrupted neuronal integrity in human neurodegenerative diseases (*78*). It will be interesting to find out whether lipid homeostasis regulated by *Hnf4* is an integral part of the *fru*-regulated neural network in pheromone detection.

Insect CHCs and pheromones undergo rapid evolution and show sexual dimorphism. In *Drosophila melanogaster*, manipulations of sex-determination genes such as *sxl*, *dsx* or *tra* lead to changed CHC profiles (*79*–*83*). Mutations or ectopic expressions of *dsx* or *tra* induces either masculinization or feminization of CHC profiles, with the major changes at sex-specific pheromones (Fig. 7B-C) (*79*, *83*). Our study revealed a novel sex-dimorphic CHC profile resulted from *fru-* or *Hnf4-* depletion in oenocytes (Fig. 3B-C, Fig. 6A-B), which differs substantially from *sxl*-, *dsx*- or *tra*-dependent sexual dimorphism of the CHC profile (*64*) (Fig. 7C-D). The male flies with *fru*- or *Hnf4*-knockdown show an almost complete loss of all CHCs, but female flies with *fru*- or *Hnf4*-knockdown have changes primarily in CHCs with the chain length > C26, which include the female pheromones 7,11-HD and 7,11-ND (Fig. 3C). The more severe changes of the male CHCs are consistent with the previous report that male CHCs change more rapidly during lab-induced natural and sexual selection in *D. simulans* (*84*). The sex dimorphic CHC patterns regulated by *dsx* or *tra* appear to stem from their regulation of DesatF and EloF expression, respectively (*24*), which promote female long-chain hydrocarbon biosynthesis (Fig. 7C) (*25*, *26*). In contrast, male flies withs *fru* or *Hnf4* knockdown have significantly reduced pheromones, and their behavioral phenotype is consistent with a previous report that wildtype males exhibit a higher preference on males without CHCs (similar to the *oeno*>*fru^IR^* generated in this study) over wild-type females (*45*). Furthermore, as we did not see changes in Fru^COM^ or HNF4 expression when *dsx* was knocked down in oenoctyes, *dsx* and *fru*/*Hnf4* likely target different downstream genes CHC and pheromone biosynthesis (Fig. 7A).

The sexual differences in hormone homeostasis may contribute to the novel sexual dimorphic CHC profiles induced by *fru*- and *Hnf4*-knockdown. CHCs need to be replenished after each molt, a process induced by pulses of ecdysteroid hormones (ecdysone). They are first stored internally before molting and subsequently transferred to the cuticular surface (*23*). It has been postulated that CHC biosynthesis is regulated by an interplay of ecdysone and juvenile hormone (JH) (*85*, *86*). As *fru* have been shown to be subject to ecdysone and JH regulation (*74*, *87*–*89*), and the expression of HNF4 is in response to hormonal pulses in a variety of insects (*90*, *91*), *fru* and *hnf4* may integrate the sex-dimorphic hormonal signals to regulate CHC biosynthesis.

## Materials and Methods

### Fly strains and genetics

Flies were maintained at 25 °C, 60% relative humidity, and 12-h light/dark cycle. Adults and larvae were reared on a standard cornmeal and yeast-based diet, unless otherwise noted. The Gal4/UAS driven RNAi crosses were cultured at 25°C until eclosion, and then incubated at 29°C for 7 days, and the same culture conditions were also used for the control group.

*promE*-*Gal4* and *promE*-*GS*-*Gal4* (*oeno*-*specific GAL4*) were obtained from Dr. H. Bai(*92*). *UAS*-*fru^MA^* and *UAS*-*fru^COMB^* were from Dr. M. Arbeitman. *elav*-*Gal4* (BDSC#6920), *UAS*-*fru^RNAi^* (BDSC#31593), *UAS*-*GFP* (BDSC#5413), *fru*-*NP21*-*Gal4* (BDSC# 30027), *fru*-*P1*-*Gal4* (BDSC#66696), *Hnf4*::*GFP.FLAG* (BDSC#38649), *UAS*-*Hnf4^RNAi^* (BDSC#29375), *UAS*-*Hnf4^RNAi^* (BDSC#64988), *UAS*-*dsx^RNAi^* (BDSC#55646), *Act5C*-*GAL4* (BDSC#4414) and *W1118* (BDSC#5905) were obtained from the Bloomington *Drosophila* Stock Center. *UAS*-*fru*-*gRNA* (VDRC#342548), *UAS*-*fru^RNAi^* (VDRC#330035), *UAS*-*fru^RNAi^* (VDRC#105005) were obtained from the Vienna *Drosophila* Resource Center. *UAS*-*Hnf4*::*HA* (F000144) was obtained from FlyORF (Zurich ORFeome Project).

RU486 (mifepristone, Fisher Scientific) was dissolved in 95% ethanol, and added to standard food at a final concentration of 100 μM for all the experiments. For activation of the GeneSwitch (GS) Gal4 driver, flies were fed on RU486 food for 7 consecutive days, unless otherwise noted.

To temporally induce *fru* MAGIC clones, early 3^rd^ instar larvae of *w*; *UAS*-*fru*-*gRNA*/*Act5C*-*Gal4*; *HS*-*Cas9*/+ were heat shocked for 30min at 37°C, and examined at day-7 after eclosion. The *HS*-*Cas9* stock was kindly provided by Dr. C. Han(*42*).

### Behavioral Assays

For the single-pair courtship assay, a tester male and a target fly (both at day-7 after eclosion) were gently aspirated into a round two-layer chamber (diameter: 1 cm; height: 3 mm per layer) to allow courtship test. Courtship index (CI), the percentage of observed time a fly performing any courtship step, was used to measure courtship between the tester male and the target fly. Each test was performed for 1 hr.

For male chaining assay, the tester males were loaded into large round chambers (diameter: 4 cm; height: 3 mm) by cold anesthesia. Chaining index, which is the percentage of observed time during which at least three flies engaged in a courtship, was used to measure the courtship behavior in groups of 13 males.

For the two-choice courtship assay, a standard courtship assay was used to test male preference (i.e., female attractiveness) between *fru* knockdown (*oeno*>*fru* ^*IR*^) and the control (*oeno*-*Gal4*) females. Assays were performed under both normal and dark conditions. For each measurement, two subject females for comparison were decapitated and placed on the opposite sides of a single well in a standard 12-well cell culture plate containing standard fly medium, and a 5-7-day old *w1118* virgin male was subsequently aspirated into the cell. The videorecording lasted for one hour to record the courtship behaviors (including orientation, wing vibration and attempted copulation) of the male fly directed toward each female. The assays were conducted at 25°C at night. To control for individual variability, male preference was shown as the percentage of time the *w1118* male was courting the *fru*-knockdown female divided by the total courtship time.

For social space assay, the tester males were loaded into large round chambers (diameter: 4 cm; height: 3 mm) by cold anesthesia. Flies were allowed to acclimate for 10 min and then digital images were collected after the flies reached a stable position (up to 20 min). Digital images were imported in Yolov5 and an automated measure of the nearest neighbor to each fly was determined using our own MAFDA system.

For food intake assay, the tester males were analyzed using a horizontal circular chamber (diameter: 4 cm; height: 3 mm). Flies were allowed to acclimate for 10 min and then digital images were collected after the flies reached a stable position (up to 20 min). Digital images were imported in Yolov5, the location of food was given manually, using our own MAFDA system to determine the number of foraging for each fly through its distribution and dwell time.

### Development of <u>m</u>achine-learning-based <u>a</u>utomatic <u>f</u>ly-behavioral <u>d</u>etection and <u>a</u>nnotation (MAFDA) system

The tracking algorithm was achieved by sorting the nearest position of fly between adjacent frames. Each fly is identified and assigned a specific ID by the system in the first frame. Subsequently, the position of each fly in the next frame is paired with the nearest neighbor in the previous frame, and inherits its ID. In the event that a target fly is lost, the fly ID remains at the position of the previous frame until a target reappears. ID-switch is identified manually. A GUI software based on Python-3.8 (Kivy-2.00) was developed to correct the IDs after ID-switching.

Yolov5 was chosen as the platform to train the data with 800 well-annotated pictures. After argumentation (flip, merge, rotate, brighten, flip, and random-salt mask), we obtained over ~140,000 labeled flies, ~130,000 labeled heads, and collected ~ 40,000 chasing events, ~8,700 wing-expansion, and ~1600 mountings. We used the Loni server to train the data with default hyperparameters with yolov5x.pt as the initial weight. The model was trained with parameters (500 epochs, 80 batch-size and 640*640 pixel) in Tesla-V100 for over a day. Chaining events are identified through 2 or more independent chase events based on a shared chasing target.

To generate an event map of fly behaviors in real-time using the MAFDA system, each fly in each frame (30 frames per second) is marked as a dot and the frames are overlayed. Different behavioral events are marked by different colored dots. To avoid over-representation, event maps are generated with 1.5-min movies to exhibit trajectories and behavioral classifications of flies in a given group.

The behavior indexes (including chasing index, singing index and chaining index) are calculated with the detected behavior events divided by the maximum possible value for that event to occur. Index = B(d)/B(max) ×100%. Chasing and singing B(max) equate fly number (N), and chaining B(max) equates N-1. For example, 10 flies in a group could have max chasing events of 10 when they form a ring and the index is 100%. When only 5 chasing-pairs are detected in a 10-fly group, the index is 50%.

### Cuticular hydrocarbon analysis

Virgin male and female flies were collected at emergence and cultured at 29°C for 7 days. Each sample (including 10 flies) was frozen at −20°C for 15 min, and then introduced to a 2 ml vial containing 100 μl hexane (Thermo Scientific) with *n*-C19 alkanes (10 ng/μl, MilliporeSigma) as an internal standard. After extraction for 5 min, 1 μl solution was injected into a GC (Trace 1310, Thermo Scientific) in splitless mode equipped with a TG-5MS capillary column (30 m × 0.25 mm × 0.25 μm, Thermo Scientific), using helium as carrier gas (1.0 ml/min). Column temperature was programmed to increase from 90°C to 150°C at 20°C/min and then to 300°C at 3°C/min. The GC was coupled with a MS (ISQ 7000, Thermo Scientific). Injection temperature was 280°C, MS source temperature was 310°C, and transfer line was 300°C. The MS was set to scan a mass range from 40 to 550. Standard mixture of *n*-alkanes (C7 to C40, MilliporeSigma) was injected following the same temperature program. The identity of 7(*Z*)-tricosene, 11-*cis* vaccenyl acetate and 7(*Z*), 11(*Z*)-heptacosadiene was confirmed by comparison of retention times and mass spectra with synthetic standards (Cayman Chemical, Cat# 9000313, 10010101 and 10012567, respectively). Other compounds were tentatively identified based on electron ionization mass spectra and Kovats indices, as well as previously published data in *Drosophila*(*93*, *94*). The quantities of CHCs were calculated based on peak areas in comparison with the internal standard. For each analysis, five biological replicates and two technical replicates were conducted.

### RNA-seq and data analysis

Adult oenocytes were carefully dissected in 1 × PBS before RNA extraction. Total RNAs were extracted from 7-day-old adult oenocytes using the Zymo RNA preparation kit. Two replicates were collected for each genotype. NEBNext Poly(A) mRNA Magnetic Isolation Module and NEBNext Ultra II RNA Library Prep Kit for Illumina were used for library preparation. The libraries were sequenced using an Illumina HiSeq 2500 system, obtaining 40 million reads for each sample. Raw reads were aligned to the *Drosophila melanogaster* reference genome (*95*). Feature Counts and DESeq2 were then used to assign gene identity to genomic features, and to analyze fold change (log2(FC)) from count data.

### cDNA Synthesis and RT-qPCR

Reverse transcription was performed on 0.25-1μg total RNA using the Superscript Reverse Transcriptase II from Thermo Fisher Scientific (Cat# 18064022) and oligo(dT) primers. cDNA was used as a template for qPCR.

qPCR was performed with a Quantstudio-3 Real-Time PCR System and PowerUp SYBR Green Master Mix (Thermo Fisher Scientific). Two independent biological replicates were performed with three technical replicates each. The mRNA abundance of each candidate gene was normalized to the expression of *Rp49* by the comparative CT methods. Primer sequences are listed in Fig.S7C.

### Generation of anti-Fru^COM^ antibody (anti-Fru^ALL^)

The rabbit polyclonal antibody against Fru^COM^ was generated by ABclonal (Wuhan, China). In brief, the fragment of the *fru* gene encoding 89 amino acids from the common part of the polypeptide, RERERERERERDRDRELSTTPVEQLSSSKRRRKNSSSNCDNSLSSSHQDR HYPQDSQANFKSSPVPKTGGSTSESEDAGGRHDSPLSMT, was cloned into the expression vector pET-28a (Sigma-Aldrich, #69864). A SUMO-tagged Fru^COM^ fusion antigen was synthesized from bacteria, purified, and used to immunize the rabbit. The anti-Fru^COM^ antibody was affinity purified. This polyclonal antibody recognizing an epitope in exon C3 of *fru*, which is present in both Fru^COM^ and Fru^M^ isoforms.

### Immunostaining and Confocal Imaging

Tissue samples were dissected in phosphate buffered saline (PBS), then fixed in 4% formaldehyde in PBS for 20 minutes. After washing with PBS with 0.2% Triton X-100 (PBT), the samples were incubated in PBT with primary antibodies at 4°C overnight with shaking and then washed in PBT three times for 15 minutes each. The following antibodies were used in immunostaining: anti-GFP (Cell Signaling Technology, #2956S, 1:200), anti-HA (Cell Signaling Technology, #3724S, 1: 500), anti-HNF4 guinea pig polyclonal antibody (1:100, a gift from G. Storelli), anti-Fru^COM^ rabbit polyclonal antibody (1:500) and rabbit anti-Fru^M^ polyclonal antibody (1:250) (*54*). The secondary antibodies conjugated with Alexa 546 or 633 (Invitrogen) were diluted 1:200 and incubated at room temperature for 2 hours. Nuclei were labeled with DAPI (Invitrogen, 1:1000). After washing, samples were mounted and imaged with Zeiss LSM 800 or Zeiss LSM 980 Confocal Microscopes. Image analysis was performed in ImageJ.

### TAG Assays

For TAG assays, 8 whole adult or 20 adipose tissues were homogenized in 100 μl PBS, 0.5% Tween 20 and immediately incubated at 70°C for 5 min. Heat-treated homogenate (20 μl) was incubated with either 20 μl PBS or Triglyceride Reagent (Sigma T2449-10ML) for 30 min at 37°C, after which the samples were centrifuged at maximum speed for 3 min. Then, 30 μl was transferred to a 96-well plate and incubated with 100 μl of Free Glycerol Reagent (Sigma F6428-40ML) for 5 min at 37°C. Samples were assayed using a BioTek Synergy HT microplate spectrophotometer at 540 nm. TAG amounts were determined by subtracting the amount of free glycerol in the PBS-treated sample from the total glycerol present in the sample treated with Triglyceride reagent. TAG levels were normalized to protein amounts in each homogenate using a Bradford assay (Bio-Rad), and data were analyzed using a student’s t test. Independent experiments were performed 2-3 times.

### Thin Layer Chromatography (TLC)

Homogenize 20 adult flies’ adipose tissue in 2:1:0.8 of methanol:chloroform:water and further using 1.4 mm ceramic beads by vigorous vortexing for 10 min at 4°C. Samples were incubated in a 37°C, water bath for 1 hour. Chloroform and 1M KCl (1:1) were added to the sample, centrifuged at 3000 rpm for 2 min, and the bottom layer containing lipids was aspirated using a syringe. Lipids were dried using argon gas and resuspended in chloroform (100 ul of chloroform/7mg of fly weight). Extracted lipids alongside serially diluted standard neutral lipids of known concentrations were separated on TLC plates using hexane:diethylether:acetic acid solvent (80:20:1). TLC plate was air dried for 10 min, spray stained with 3% copper (II) acetate in 8% phosphoric acid and incubated at 180°C in the oven for 10 min to allow bands to develop for scanning and imaging.

### Starvation

Polystyrene vials (Genesee Scientific Cat# 32-110) were half filled with water and a dense weave cellulose acetate stopper (Genesee Scientific Cat# 49-101) was pushed to the bottom of the vial allowing saturation of this artificial substrate with water. Excess water was discarded and newly emerged flies were transferred to these vials at a density of 5-10 males per vial. Vials were sealed with a second dense weave cellulose acetate stopper to reduce water evaporation. Starvation experiments were conducted at 29°C. In this case, animals were transferred daily to fresh starvation vials and collected at 6-7 days after emergence for lipid stains.

### Lipid Stains

Oenocytes dissected from adults were fixed in 4% paraformaldehyde for 30 min at room temperature, rinsed in PBS, and incubated in Bodipy 493/503 (1:1000, Thermo Fisher Scientific Cat#D3922) or NileRed (1:2000, TCI America, 7385-67-3) for 1hr at room temperature, in the dark, to stain the neutral lipid droplets. Next, the oenocytes tissues were rinsed in PBS and mounted in antifade reagent with DAPI (Invitrogen).

### Quantification and Statistical Analysis

Data analyses were conducted in GraphPad Prism. Unpaired t-test was used for two-sample comparisons, one-way ANOVA Dunnett’s multiple comparison test and One-way ANOVA with Tukey’s multiple comparison test were used for multiple-sample comparisons. Specific statistical approaches for each figure are indicated in the figure legend.

## Acknowledgements

We thank Dr. M. Arbeitman, H. Bai, C.-Y. Lee, C. Han, Bloomington *Drosophila* Stock Center (BDSC, USA) and the Vienna Drosophila RNAi Center (VDRC, Austria) for providing fly stocks; Dr. G. Storelli and the Developmental Studies Hybridoma Bank (DSHB, USA) for providing antibodies. We also thank Dr. X.-M. Yu and Dr. S. Lenhert for critical reading and helpful comments on the manuscript and members of the Deng lab for feedback and suggestions. Special thanks to A. Brown from the FSU Department of Biological Science for help with bulk RNA-seq library preparation.

## Authors’ contribution

Conception and design: J. Sun, W.-M. Deng

Development of methodology: J. Sun, W.-M. Deng

Acquisition of data (provided animals, acquired and managed animals, provided facilities, etc.): J. Sun, C. Ellsworth, Q. Sun, Y-C. Huang

Analysis and interpretation of data (e.g., statistical analysis, biostatistics, computational analysis): J. Sun, W.-K. Liu

Study supervision: J. Sun, W.-M. Deng

Writing – original draft: J. Sun, W.-M. Deng

Writing – review & editing: J. Sun, W.-M. Deng, Q. Sun, Y.-F. Pan

Acquisition of funding – W.-M. Deng

All authors read and approved the final manuscript.

## Data and materials Availability

All data used to evaluate the conclusions are presented in the paper and/or the Supplementary Materials. Requests for further information and additional data and materials related to this paper should be directed to and will be fulfilled by the corresponding author, Wu-Min Deng.

