## Supplemental Figure S1 to S7 for "Integrating lipid metabolism, pheromone production and perception by Fruitless and Hepatocyte nuclear factor 4"

**This file includes:**

Figs. S1 to S7.

Movie. S1 to S5.

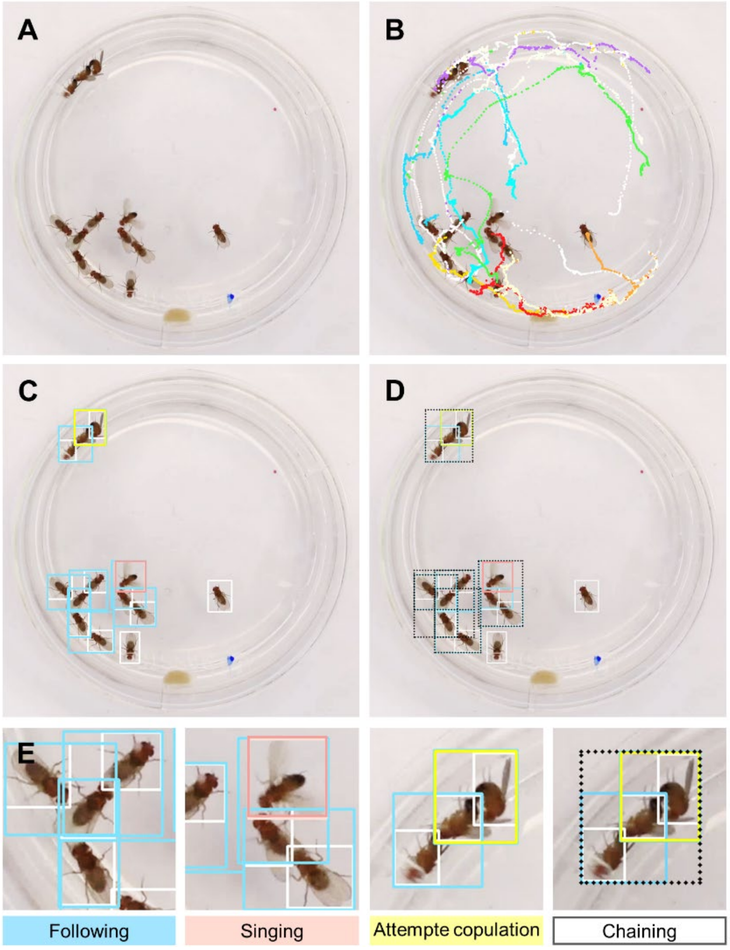


**Fig. S1. The machine-learning-based automatic fly-behavioral detection and annotation (MAFDA) system. (**A) A snapshot of the original video. (B) The tracked trajectory of individual flies indicated by different colored lines. (C-D) Identification of different behavior types for auto-annotation (head box not shown here). (E) Different colors represent different behavior types. White boxes mark flies in resting/walking. Blue boxes mark flies that are following. Salmon box shows singing flies. Yellow box shows flies in copulation. Black dashed boxes mark chaining flies.

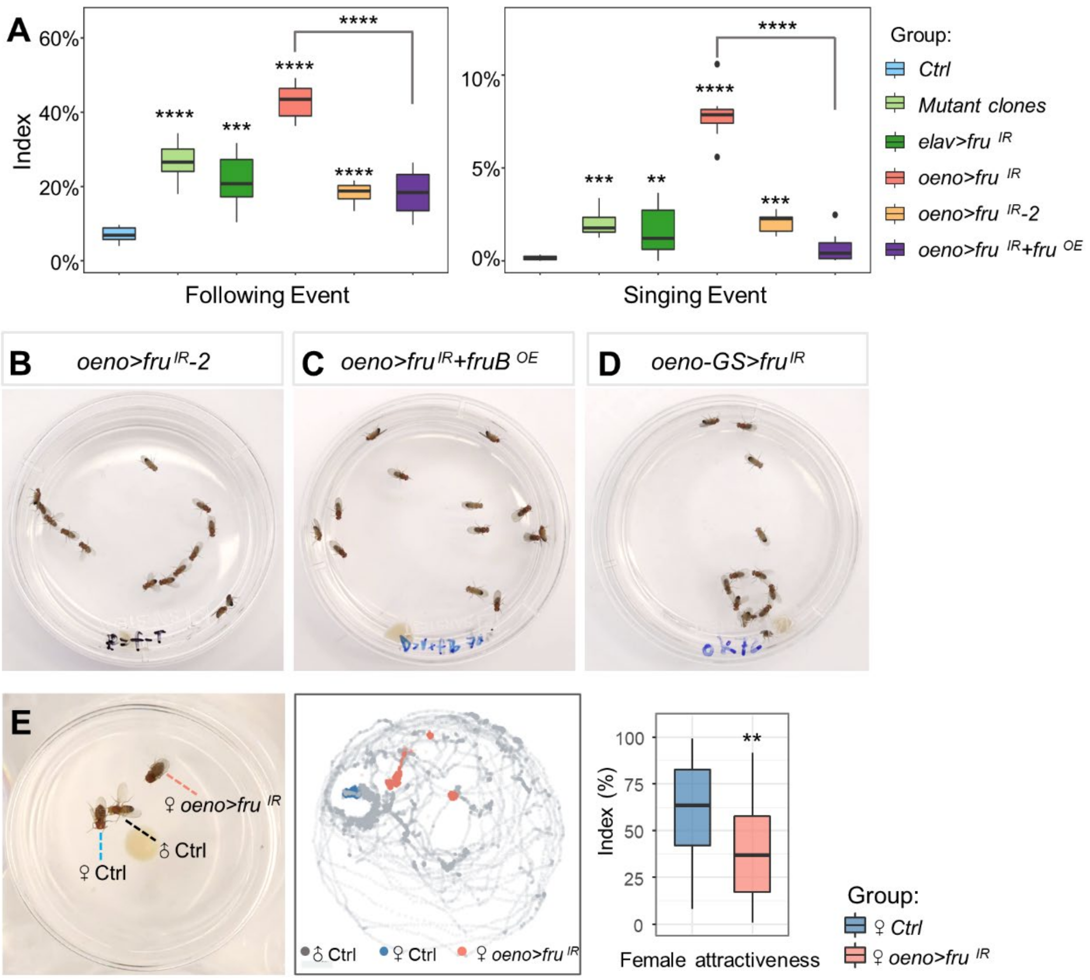


**Fig. S2. Fly behavioral phenotyping of *fru* knockdowns in oenocytes.** (A) Quantitative statistical results of following and singing index for Fig.1A. There were 13 flies per video with 8 independent biological replicates. (B-D) Representative video screenshots of each fly group with indicated genotype. (B) Male flies with knockdown of *fru* using an alternative *RNAi* (BDSC#31593) in oenocytes also exhibits the male-male chaining behavior (C) Fru$^{COMB}$ overexpression alleviates the defective behavior caused by *fru* knockdown in the oenocyte. (D) Using an oenocyte-specific GeneSwitch driver (*promE-GS-Gal4*), knockdown of *fru* in adult oenocytes induced male-male chaining behavior. (E) Oenocyte-specific *fru*-depletion in female flies reduced their sexual attractiveness to males. An event map generated from a 1-hour video of the two-choice courtship assay. Grey dots indicate the *control* (*oeno/+*) male, blue dots indicate the headless *control* (*oeno/+*) female. Salmon dots indicate the headless *oeno>fru*$^{IR}$ female. Boxplots show that *oeno>fru*$^{IR}$ females have lower courtship index (CI) than its heterozygote parental control lines (n=36), as revealed in the two-choice courtship assay. All data are represented as mean ± SEM. P values are calculated using one-way ANOVA followed by Holm-Sidak multiple comparisons. **p<0.01, ***p<0.001.

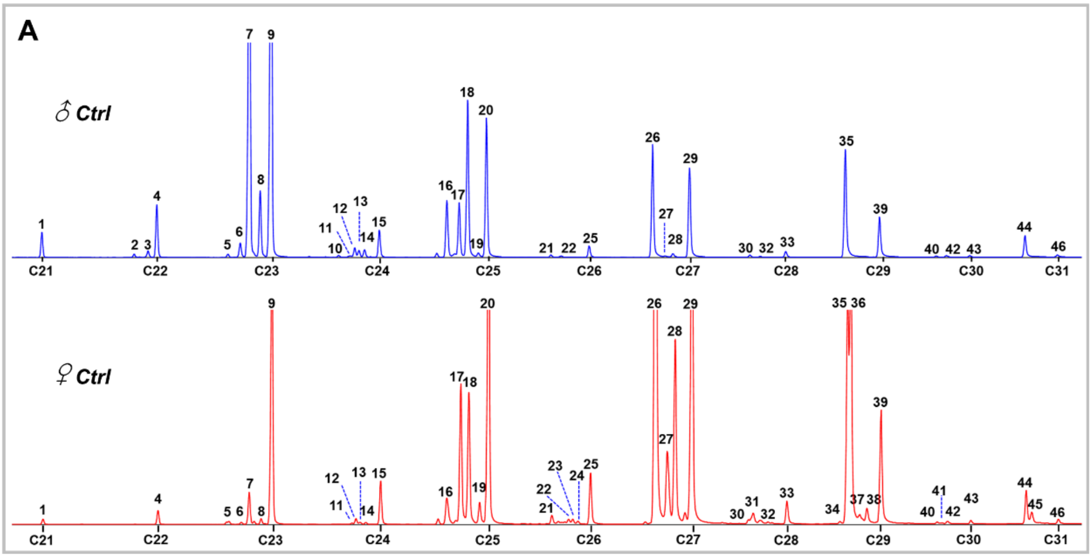


**B**

| No. | Retention Time (min) | Compound | Chemical Class | Kovats index | No. | Retention Time (min) | Compound | Chemical Class | Kovats index |
|---|---|---|---|---|---|---|---|---|---|
| 1 | 22 | n-Heneicosane | alkane | 2100 | 23 | 34.628 | (Z)-9-Hexacosene | monoene | 2582 |
| 2 | 24.203 | (+)-6-Docosene | monoene | 2180 | 24 | 34.753 | (Z)-7-Hexacosene | monoene | 2588 |
| 3 | 24.530 | cis-vaccenyl acetate | cVA | 2192 | 25 | 35.053 | n-Hexacosane | alkane | 2600 |
| 4 | 24.741 | n-Docosane | alkane | 2200 | 26 | 36.553 | 7,11-C27:2 (7,11-HD) / 2-Me-C27 | diene / Me-alkane | 2663 |
| 5 | 26.438 | 7,11-C23:2 (7,11-TD) | diene | 2262 | 27 | 36.849 | (Z)-9-Heptacosene | monoene | 2675 |
| 6 | 26.727 | (Z)-9-Tricosene | monoene | 2273 | 28 | 37.036 | (Z)-7-Heptacosene | monoene | 2683 |
| 7 | 26.962 | (+)-7-Tricosene | monoene | 2281 | 29 | 37.437 | n-Heptacosane | alkane | 2700 |
| 8 | 27.206 | (+)-5-Tricosene | monoene | 2290 | 30 | 38.818 | 7,11-C28:2 (7,11-OD) | diene | 2760 |
| 9 | 27.475 | n-Tricosane | alkane | 2300 | 31 | 38.927 | 2-Me-C27 | Me-alkane | 2765 |
| 10 | 29.074 | x,y-C24:2 | diene | 2362 | 32 | 39.090 | Octacosene | monoene | 2774 |
| 11 | 29.332 | 3-Me-C23 | Me-alkane | 2372 | 33 | 39.733 | n-Octacosane | alkane | 2800 |
| 12 | 29.455 | (+)-7-Tetracosene | monoene | 2377 | 34 | 40.998 | 9,13-C29:2 | diene | 2856 |
| 13 | 29.557 | (+)-6-Tetracosene | monoene | 2381 | 35 | 41.175 | 2-Me-C28 | Me-alkane | 2864 |
| 14 | 29.689 | (Z)-5-Tetracosene | monoene | 2386 | 36 | 41.256 | 7,11-C29:2 (7,11-ND) | diene | 2868 |
| 15 | 30.043 | n-Tetracosane | alkane | 2400 | 37 | 41.467 | (Z)-9-Nonacosene | monoene | 2877 |
| 16 | 31.652 | 7,11-C25:2 (7,11-PD) | diene | 2463 | 38 | 41.644 | (Z)-7-Nonacosene | monoene | 2885 |
| 17 | 31.948 | (x)-9-Pentacosene | monoene | 2474 | 39 | 41.981 | n-Nonacosane | alkane | 2900 |
| 18 | 32.158 | (Z)-7-Pentacosene | monoene | 2483 | 41 | 43.416 | 2-Me-C29 | Me-alkane | 2967 |
| 19 | 32.403 | (x)-5-Pentacosene | monoene | 2492 | 43 | 44.113 | n-triacontane | alkane | 3000 |
| 20 | 32.601 | n-Pentacosane | alkane | 2500 | 44 | 45.426 | 2-Me-C30 | Me-alkane | 3063 |
| 21 | 34.131 | 2-Me-C25 | Me-alkane | 2563 | 45 | 45.566 | 7,11-C31:2 | diene | 3070 |
| 22 | 34.532 | 7,11-C26:2 | diene | 2574 | 46 | 46.198 | n-Hentriacontane | alkane | 3100 |

**Fig. S3. GC-MS analysis of cuticular hydrocarbon extracts from control males and females.** (A) Sexually dimorphic CHC profiles in *Drosophila melanogaster*. The graphs show representative chromatograms of CHCs of 7-day-old virgin male and female flies, with the male at the top (blue) and female at the bottom (red). Compounds corresponding to each numbered peak are listed in B. Compounds that are shared between sexes bear the same number. The identity of each hydrocarbon was confirmed by comparison with synthetic standards and Kovats index in publication *(1, 2)*.

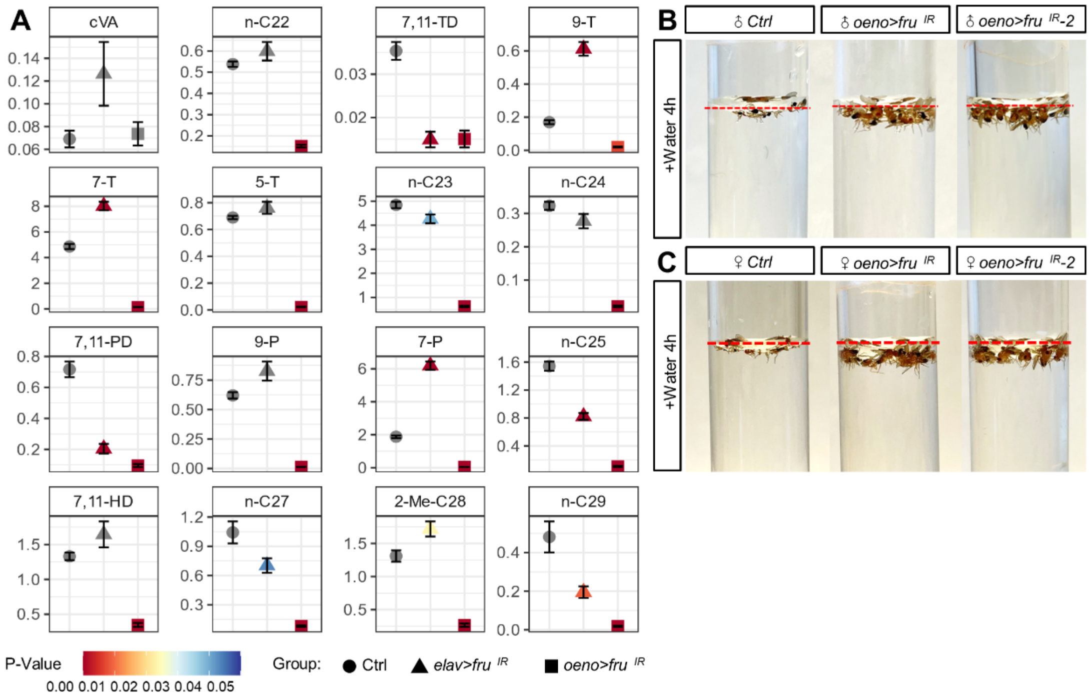


**Fig. S4. Oenocyte-specific knockdown of *fru* decreased representative cuticular hydrocarbons and waterproof capacity.** (A) 16 representative cuticular hydrocarbons were selected for quantitative statistics and normalized by the internal standards. 5 independent biological replicates were collected for each genotype. Circles indicate the *control* group, triangles indicate the *elav>fru*$^{IR}$ group and squares indicate the *oeno>fru*$^{IR}$ group. Different p-values are shown by gradient colors. (B and C) The cuticular hydrophobicity of control, *oeno>fru*$^{IR}$ and *oeno>fru*$^{IR}$*-2* was tested following death by dry starvation and incubation of the carcasses in water with agitation. After 4 hours, dead control carcasses continue to float above the liquid (represented by the red dotted line), while *oeno>fru*$^{IR}$ and *oeno>fru*$^{IR}$*-2* equilibrate below the surface (n=13 males in each vial). Males are shown in a and females in b.

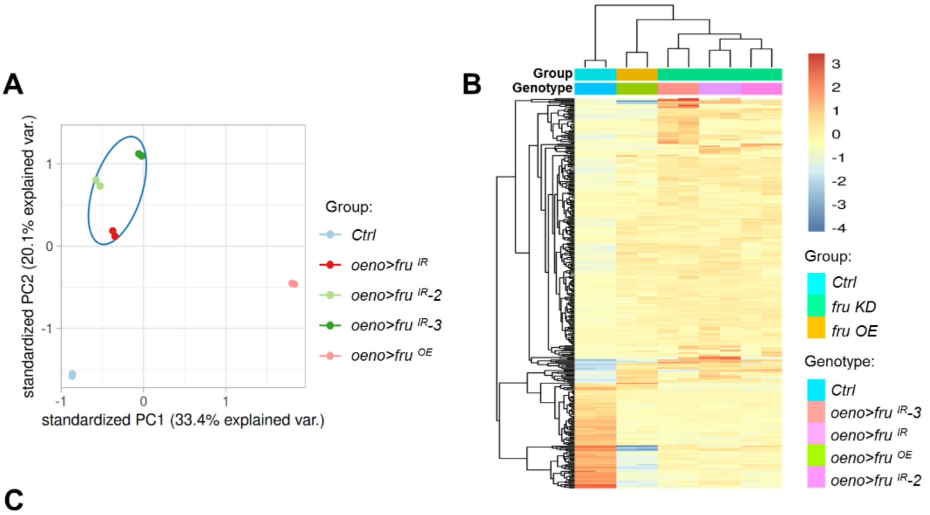


**C**

| Name | Function | Forward primer | Reverse primer |
|---|---|---|---|
| ACC | Acetyl-CoA carboxylase | ATGAGCGAAACAAATGAGTCCA | GGAACTCGTCACATGCCTCG |
| Hnf4 | Fatty acid beta-oxidation | GGCGACGGGCAAACATTATG | CGCAAATCTGCAAGTGTACTGAT |
| FASN1 | Fatty acid synthase | GAGCAACAACAGCCCACTTG | AGAGGCCATGAAACGGAAGG |
| FASN2 | Fatty acid synthase | CTGGACGATCCCTTCTACGC | TTACCGGAGGCCAACATCAC |
| desat1 | Fatty acid desaturase | TGCCGATTGCTTGCTTCAT | TTCACCCCAGGCGTACATG |
| desat2 | Fatty acid desaturase | GGTGGTGCTTCCAGCTAAACA | GGCGATTTCCGAATTTATGG |
| CG16904 | Elongase | GAGGTGTGCTCAAGTTCTACAAC | TGCACGAATAACAGATGGATGC |
| CG30008 | Elongase | ATCCAGGTGGTTTCCTGCATA | GCGGCACTTCCAAAAGATGT |
| Cyp4g1 | Aldehyde decarbonylase | TGGAAGTAGTTCAGGAGACGC | GGTGCCCACTAAGGTGGTT |
| Cpr | Cytochrome P450 reductase | GCAGCCTGACCGCATTTAC | AGGAATGGGTTCTTGGCGTC |
| rp49 | Internal reference | GACGCTTCAAGGGACAGTATCTG | AAACGCGGTTCTGCATGA |

**Fig. S5. Bulk RNA-seq analysis to identify genes affected by *fru* knockdown in oenocytes.** (A) PCA plot shows that the biological replicates of each genotype are highly consistent. The datasets from *fru*-knockdown induced by three independent RNAi lines show good clustering. (B) Heat map of expression of differentially expressed genes identified from bulk RNA-seq. Gene expression is shown in normalized log2 counts per million. Differentially expressed genes were selected based on a fourfold change and FDR<0.05. (C) Primer sequences for RT-qPCR of selected genes involved in VLCFA/hydrocarbon biosynthesis.

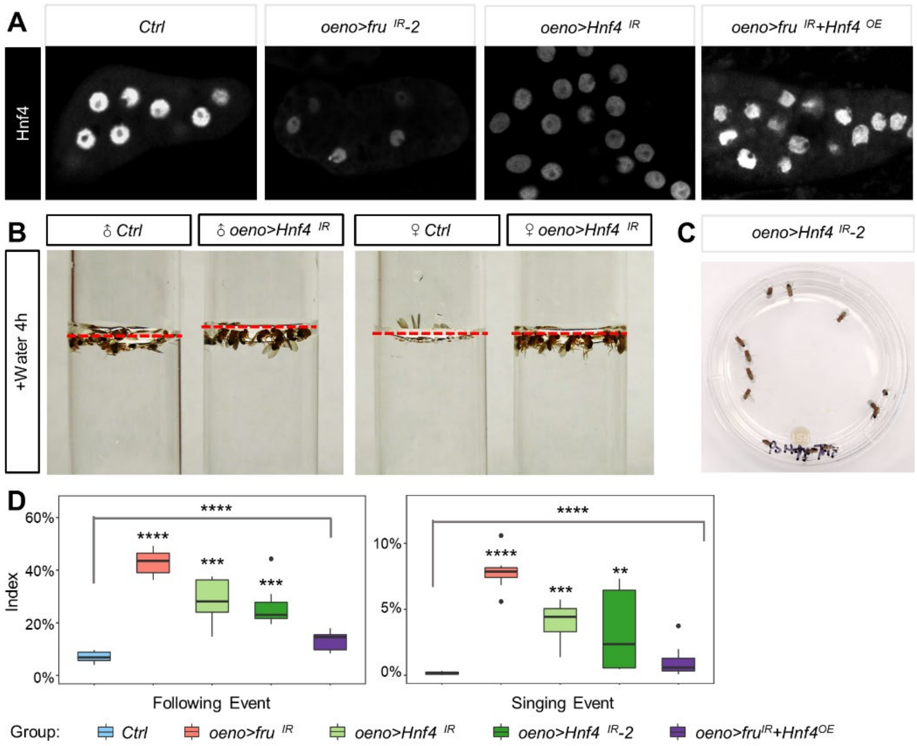


**Fig. S6. *Hnf4* loss in oenocytes causes behavioral and CHC synthesis defects.** (A) Using the anti-HNF4 antibody to detect the protein level of HNF4 in *oeno>fru*$^{IR}$*-2, oeno>Hnf4*$^{IR}$ and *oeno>fru*$^{IR}$*+Hnf4*$^{OE}$. (B) The cuticular hydrophobicity of control and *oeno>Hnf4*$^{IR}$ was tested following death by dry starvation and incubation of the carcasses in water with agitation. After 4 hours, dead control carcasses continue to float above the liquid (represented by the red dotted line), while *oeno>Hnf4*$^{IR}$ equilibrate below the surface in males and females (n=13 files in each vial). (C) A representative video screenshot of group-housed *oeno>Hnf4*$^{IR}$*-2* flies. Knockdown of *Hnf4* in oenocytes with a different *Hnf4 RNAi* (BDSC#64988) also showed male-male chaining behavior. (D) Quantitative statistical results of the following and singing behaviors for Fig.4A. There were 13 flies per video with 8 independent biological replicates. Data are represented as mean ± SEM. P values are calculated using one-way ANOVA followed by Holm-Sidak multiple comparisons. ***p<0.001.

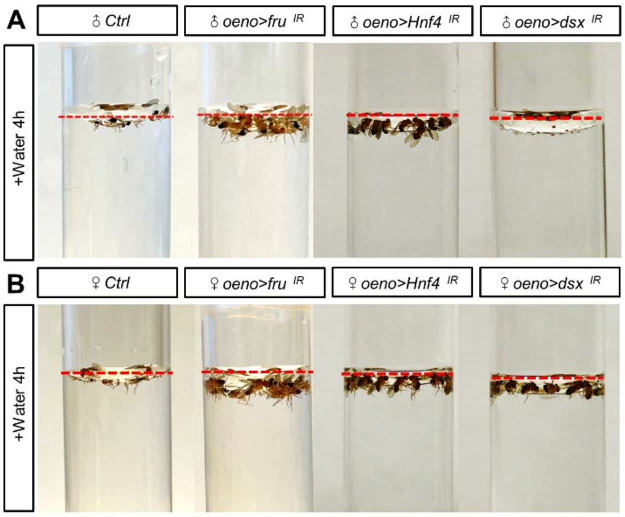


**Fig. S7. Deletion of *dsx* results in sexually dimorphic water resistance.** (A and B) The cuticular hydrophobicity of *oeno*>*dsx*$^{IR}$ was tested following death by dry starvation and incubation of the carcasses in water with agitation. After 4 hours, dead *oeno*>*dsx*$^{IR}$ males carcasses continue to float above the liquid (represented by the red dotted line), while *oeno*>*dsx*$^{IR}$ females equilibrate below the surface (n=13 males in each vial).
